# Nucleosome conformation dictates the histone code

**DOI:** 10.1101/2022.02.21.481373

**Authors:** Matthew R. Marunde, Harrison A. Fuchs, Jonathan M. Burg, Irina K. Popova, Anup Vaidya, Nathan W. Hall, Matthew J. Meiners, Rachel Watson, Sarah A. Howard, Katherine Novitzky, Eileen McAnarney, Marcus A. Cheek, Zu-Wen Sun, Bryan J. Venters, Michael-C. Keogh, Catherine A. Musselman

## Abstract

Histone post-translational modifications (PTMs) play a critical role in chromatin regulation. It has been proposed that these PTMs form localized ‘codes’ that are read by specialized regions (reader domains) in chromatin associated proteins (CAPs) to regulate downstream function. Substantial effort has been made to define [CAP-histone PTM] specificity, and thus decipher the histone code / guide epigenetic therapies. However, this has largely been done using a reductive approach of isolated reader domains and histone peptides, with the assumption that PTM readout is unaffected by any higher order factors. Here we show that CAP-histone PTM interaction is in fact dependent on nucleosome context. Our results indicate this is due to histone tail accessibility and the associated impact on binding potential of reader domains. We further demonstrate that the *in vitro* specificity of a tandem reader for PTM-defined nucleosomes is recapitulated in a cellular context. This necessitates we refine the ‘histone code’ concept and interrogate it at the nucleosome level.

---

The eukaryotic genome exists in the cell nucleus in the form of chromatin, a complex between DNA and histone proteins. The basic repeating chromatin subunit is the nucleosome (hereafter Nuc): an octamer of core histones (two each of H2A, H2B, H3, and H4) wrapped by ∼147 base pairs of DNA (**Fig. 1a**)^1^. Chromatin organization is critical for regulation of the underlying genome and is spatially and temporally controlled throughout development and within somatic cells. One of the major mechanisms to modulate chromatin structure is post-translational modification (PTM) of the histone proteins, particularly on their N-terminal and C-terminal tails (**Fig. 1a**). Globally speaking, particular histone PTMs are correlated with distinct chromatin states (*e*.*g*. transcriptional activation/repression, damaged DNA) and genomic elements (*e*.*g*. gene enhancers, centromeres)^2–4^. Importantly, it has been proposed that histone PTMs function in diverse combinations, perhaps even forming a ‘histone code’^5–7^ read by <u>C</u>hromatin-<u>A</u>ssociated <u>P</u>roteins (CAPs) via their various ‘reader domains’, thus localizing and/or regulating CAP activity^8,9^. However, the dictates of such a code and role that reader domains play in mediating same is hotly debated, as it has been challenging to: determine the pattern of PTMs read out by tandem domains *in vitro*, determine whether this pattern is actually being discerned in the *in vivo* context, and finally determine if this leads to a unique biological outcome^10^. Resolving this situation is critical not only to define the fundamentals of the histone code, but also as efforts ramp up to utilize PTM patterns in disease diagnostics and therapeutically target CAP-PTM associations^11–14^.

**Fig. 1.**
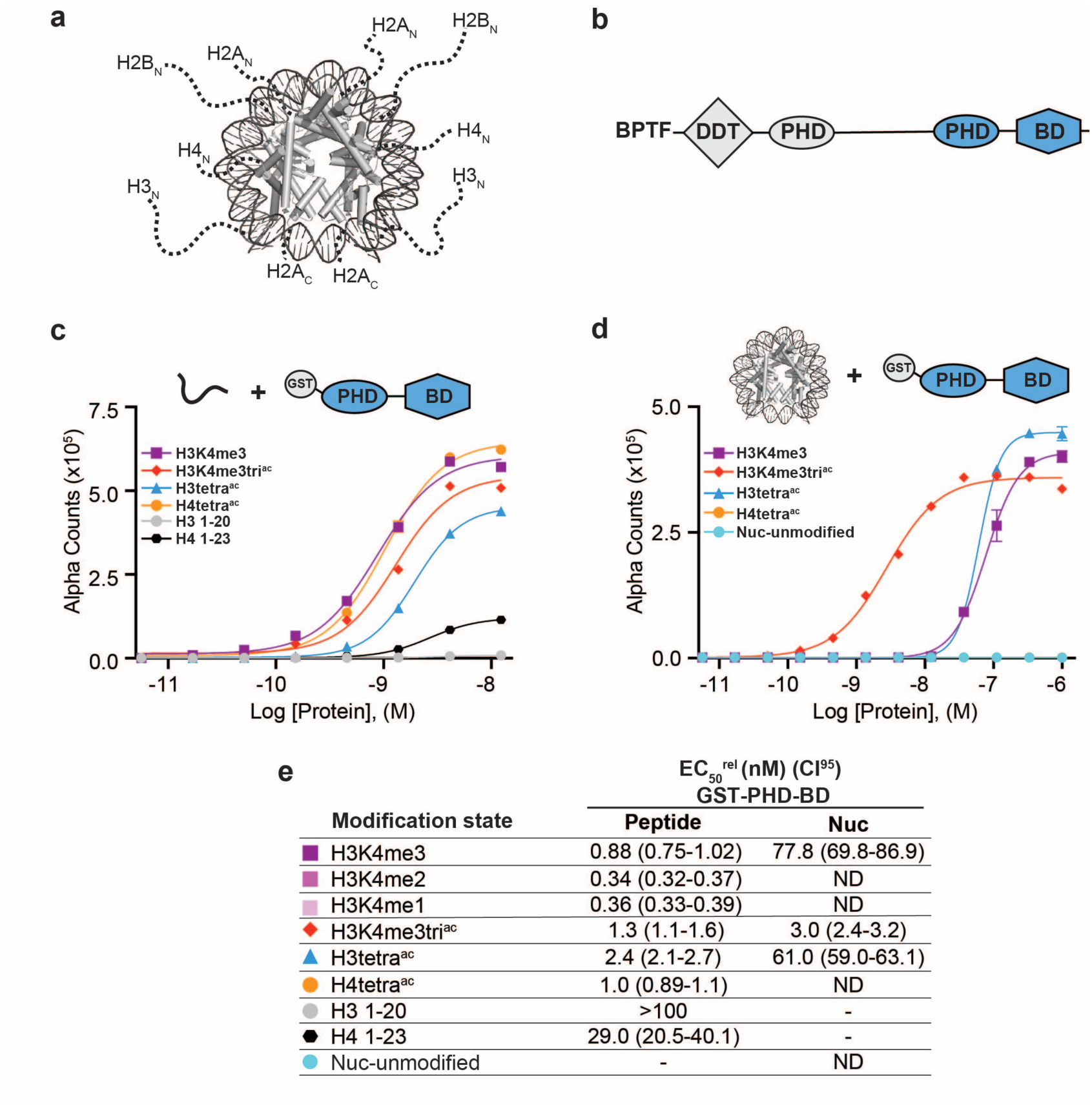
BPTF PHD-BD demonstrates restricted and synergistic PTM binding in the nucleosome *vs*. peptide context. **a)** The nucleosome core particle (PDB: 3LZ0) (Nuc): histone N- and C-terminal tails (as defined by trypsin digest and to relative scale) are depicted as dotted lines. **b)** Secondary domain architecture of BPTF [Uniprot Q12830; 3,046 aa; 338 kDa]. Region covered by the C-terminal tandem PHD-BD (aa 2865-3036; as used throughout this study) is in blue. **c**,**d)** Alpha counts plotted as a function of GST-PHD-BD Query concentration from dCypher assays with histone peptide (**c**) or Nuc (**d**) Targets. **e)** Relative EC_50_ (EC_50_^rel^) and 95% confidence interval (CI^95^) values from dCypher curves (in **c**,**d** and **Extended Data Fig. 1d**,**e**; for calculation see **Methods** and **Suppl. Discussion**). Targets are color coded as per legends. ND, Not Determined.

As a starting point, it is critical to clearly establish the PTM patterns actually engaged by reader domains. To date, the *in vitro* specificity of individual readers has almost exclusively been determined using modified histone tail peptides^9,15,16^. However, this approach provides no insight as to any effect of Nuc context. Moreover, our understanding of the selectivity for PTM patterns can only be inferred as a simple sum of the specificity for individual reader domains, again ignoring any effect of Nuc context^15,17^. Many enzymes that act on the histone tails (*e*.*g*. lysine demethylases / methyltransferases) show altered activity when comparing peptide and Nuc substrates^18–22^, suggesting the importance of the higher order environment. Similarly, several studies indicate that reader domains have altered affinity for histone tails in the Nuc context^23–26^. Thus, an essential question arises; does Nuc context alter histone PTM pattern readout? Here we show this is indeed the case, with not only affinity but also specificity impacted for each individual and a tandem of reader domains. We find an overall decrease in the affinity of each reader for histone tails, and a change in their preferred PTM pattern on Nucs versus peptides. Our results support this is largely due to accessibility in the Nuc context, in which histone tails must be displaced from DNA to enable PTM readout. This alters the engagement of individual domains and the multivalent activity of the tandem domains. We propose that the ‘histone code’ is ultimately defined by a combination of three elements: 1) the PTMs that can be recognized and bound by individual reader domains; 2) accessibility of the modified histone tails in the Nuc context; and, 3) the organization and multivalent binding potential of grouped domains (where the whole is greater than the sum of the parts).

## BPTF PHD-BD demonstrates restricted specificity and synergistic binding in the Nuc context

The BPTF subunit is important for chromatin association of the NURF (<u>Nu</u>cleosome <u>R</u>emodeling <u>F</u>actor) complex^27,28^, and is a pro-tumorigenic factor in several malignancies^29^. At the BPTF C-terminus is a tandem of reader domains: a PHD finger and bromodomain (PHD-BD, **Fig. 1b**). These are of interest for targeted therapeutics^30^, and so an understanding of their function is critical. In the context of histone peptides, the PHD finger (PHD) has been shown to associate with H3 tri-methylated at lysine 4 (H3K4me3)^28^, while the bromodomain (BD) binds to histone tails containing acetylated lysines, with a preference for the acetylated H4 tail^31–33^. While some efforts have been made to investigate the recruitment of BPTF PHD-BD to modified Nucs, only a limited subset of H3K4me3/H4Kac combinations based on peptide data have been tested, suggesting a preference for [H3K4me3 / H4K5acK8acK12acK16ac (hereafter H4tetra^ac^)]^33^ or [H3K4me3 / H4K16ac]^33,34^.

To more comprehensively investigate if Nuc context alters the BPTF PHD-BD readout of histone PTMs, we utilized the *dCypher***®** approach^35^ (dCypher for brevity) on the Alpha**®** platform^36,37^ (**Extended Data Fig. 1a** and **Suppl. Discussion**) to screen GST- and 6His-tagged forms of the tandem reader (GST-PHD-BD and 6His-PHD-BD; see **Suppl. Discussion**) against large panels of biotinylated PTM-defined peptides (287×) and Nucs (59 ×) (**Methods** and **Suppl. Tables 3: Resources A-C**). This no-wash bead-based proximity assay allows measurement of the relative EC_50_ (EC_50_^rel^) between Queries : Targets (*i*.*e*. readers : histone PTMs) by plotting Alpha Counts (fluorescence) as a function of protein concentration^35^ (see **Suppl. Tables 1 & 2** for all EC_50_^rel^ from this study).

In agreement with previous studies, the GST-PHD-BD Query showed strong selectivity for methylated H3K4 peptides over all other methyl-residues represented (me1-2-3 at H3K9, H3K27, H3K36, and H4K20: **Extended Data Fig. 1b**). Also in agreement with previous data, GST-PHD-BD demonstrated a preference for acetylated H4 tail peptides (**Fig. 1c,e** and **Extended Data Fig. 1b**), although we observed little difference in binding to a multiply acetylated H4 tail versus any of the singly acetylated residues (**Extended Data Fig. 1b**). We also observed comparable binding to singly or multiply acetylated H3 tail peptides, though with approximately two-fold weaker EC_50_^rel^ as compared to H4 (**Fig. 1c,e** and **Extended Data Fig. 1b**). Similar results to these targets were obtained with a 6His-PHD-BD Query (see **Suppl. Discussion**). Finally, we observed no preference for a H3K4me3K9acK14acK18ac (hereafter H3K4me3tri^ac^) peptide over those containing each PTM class alone, in agreement with a recent study^38^ (**Fig. 1c,e)**. Thus, peptides provide no support for a ‘histone code’ model, in which multivalent engagement by PHD-BD with a cognate combinatorially modified substrate would be expected to manifest as stronger binding than engagement of either domain alone.

We next examined interaction of the GST-PHD-BD Query with PTM-defined Nucs and found several striking differences. First, the overall affinity for Nucs was reduced relative to peptides (**Fig. 1c-e**). Second, binding of GST-PHD-BD to Nucs recapitulated only a subset of the interactions observed with peptides (**Fig. 1c-e** and **Extended Data Fig. 1b**,**c**). The differences included a selectivity for H3K4me3 over the me2 / me1 states (**Extended Data Fig. 1d**,**e**), and binding to acetylated H3 but not acetylated H4 (**Fig. 1d,e** and **Extended Data Fig. 1b**,**c**). A third contrast to peptides was a dramatic increase in the GST-PHD-BD affinity for Nucs containing the H3K4me3tri^ac^ combinatorial pattern versus those containing each PTM class alone (26-fold over H3K4me3; 20-fold over H3K4acK9acK14acK18ac (hereafter H3tetra^ac^)) (**Fig. 1d,e**). This last point would support a ‘histone code’ where reader domains act synergistically to engage preferred PTM patterns.

To further refine the PTM patterns recognized by GST-PHD-BD in the Nuc context, we explored additional H3 methyl / acetyl combinations (**Extended Data Fig. 1f**,**g**). Here the Query bound with similar EC_50_^rel^ to H3K4me3tri^ac^, H3K4me3K14ac, and H3K4me3K18ac, but substantially weaker to H3K4me3K9ac. Notably, crystal structures of BPTF BD in complex with acetylated histone peptides^33^ indicate that the bromodomain binding pocket can only accommodate one acetyl-lysine. Thus, data supports that in the Nuc context, PHD-BD preferentially reads out H3K4me3K14ac or H3K4me3K18ac.

### Individual domains have reduced affinity and altered specificity in the Nuc context

To gain further insight into the contribution of each domain to the synergistic binding of PHD-BD, we tested their individual reader ability for peptides and Nucs. As for the tandem PHD-BD Query, the relative affinity of 6His-PHD was reduced for Nucs relative to peptides (**Extended Data Fig. 2a-d** and **Suppl. Tables 1 & 2**). Interestingly, while on peptides 6His-PHD preferentially associated with H3K4me3 with approximately two-fold weaker EC_50_^rel^for H3K4me3tri^ac^ (**Extended Data Fig. 2a**,**c**), this relationship was inverted for Nucs (**Extended Data Fig. 2b**,**d**). The same general affinity trends were observed for the GST-PHD Query (**Extended Data Fig. 2b**,**e** and **Suppl. Discussion**). Of particular note, in testing methyl / single acetyl combinatorial Nucs GST-PHD had stronger binding with a co-incident acetyl present, but demonstrated a similar EC_50_^rel^ for K9ac, K14ac or K18ac (**Extended Data Fig. 2b**,**e**). This was a striking contrast to the interaction of tandem GST-PHD-BD with Nucs, where H3K9ac did not contribute to an improved EC_50_^rel^. In agreement with previous studies^33^, 6His-BD alone bound both acetylated H3 and H4 peptides, but with a preference for acetylated H4 (**Extended Data Fig. 2a**,**f**). However when presented with Nucs, 6His-BD showed no detectable binding to any tested targets (**Extended Data Fig. 2b**,**g**).

From above, Nuc context has a substantial (and unexpected) impact on the interaction of the individual PHD and BD with modified histone tails. PHD alone requires the combination of H3K4 methylation and H3 tail acetylation (K9ac, K14ac, K18ac), but does not discern between individual acetylated residues (see **Discussion**). BD alone cannot associate with any acetylated Nuc substrates, but requires the endogenously partnered PHD to engage the H3 tail independent of its methylation. However, the presence of BD in the tandem construct confers specificity for H3K14ac and H3K18ac **(Extended Data Fig. 1g)**.

### Both domains are required for full activity of the tandem module

To further investigate the contribution of each domain to binding in the tandem context, we created individual loss-of-function mutants of either PHD (aromatic cage W2891A; PHD^mut^) or BD (ZA-loop N3007A; BD^mut^) (**Suppl. Tables 3: Resources A**) for dCypher testing. Mutation of W2891 has no impact on PHD structure but abrogates binding to H3K4me3 peptides^27,28^; mutation of N3007 has no impact on the BD fold but dramatically reduces binding to acetylated peptides (**Fig. 2a** and **Extended Data Fig. 3**). Of note, GST-PHD^mut^-BD lost binding to all tested Nucs (H3K4me3, H3tetra^ac^, or H3K4me3tri^ac^). This is consistent with the individual domain activity and revealed that even in the tandem context an active BD is insufficient to mediate Nuc binding without a functional PHD (**Fig. 2b,c**). In contrast, GST-PHD-BD^mut^ retained Nuc engagement if provided methylated H3K4: *i*.*e*. binding H3K4me3 and H3K4me3tri^ac^ but not H3tetra^ac^ (**Fig. 2b,c**). Together this indicates that the PHD is critical for association with all Nuc targets, even when H3K4 is not methylated. In contrast BD contributes to Nuc binding only when H3 is acetylated.

**Fig. 2.**
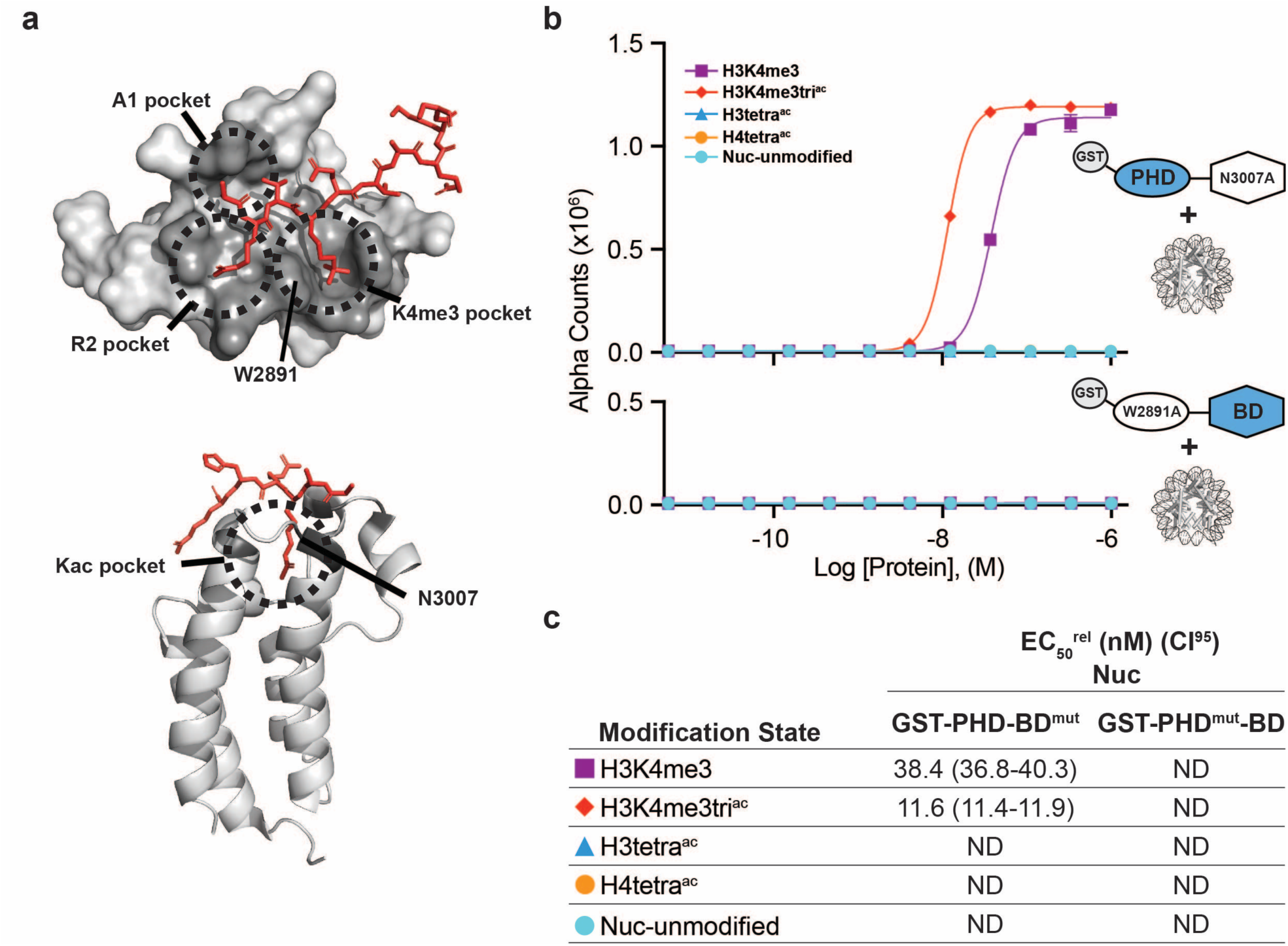
BPTF PHD and BD both contribute to Nuc binding. **a)** The PHD-H3K4me3 (top) and BD-Kac (bottom) binding pockets are highlighted on previously solved structures of the individual domains in complex with histone peptides (PDB: 2FUU and 3QZT). Binding pockets are circled / labeled: on PHD for A1, R2, and K4me3; on BD for Kac. Relative location of PTM-binding residues W2891 (PHD) and N3007 (BD) are also indicated (and mutated to alanine in **b, c**). **b)** Alpha Counts from dCypher assays plotted as a function of GST-PHD-BD^N3007A^ (GST-PHD-BD^mut^; top) or GST-PHD^W2891A^-BD (GST-PHD^mut^-BD; bottom) Query concentration to Nuc Targets. **c)** EC_50_^rel^ (CI^95^) values from dCypher curves in (**b**). Targets are color coded as per legends. ND, Not Determined.

### The PHD-BD makes multivalent contacts with the acetylated H3 tail

Above results suggest that both reader domains support the association of the tandem PHD-BD with the acetylated H3 tail independent of methylation state, whereas the BD may only associate with the acetylated H4 tail in a peptide format. To further investigate this, we turned to NMR spectroscopy. Sequential ^1^H,^15^N-HSQC spectra were recorded on ^15^N-labeled PHD-BD upon addition of unlabeled histone peptides corresponding to H3tri^ac^, H3tetra^ac^, or H3K4me3tri^ac^ (**Fig. 3a** and **Extended Data Fig. 4**). Chemical shift perturbations (CSPs) were observed in BD resonances upon addition of all three peptides, indicating ligand engagement. The bound state chemical shift for BD resonances was similar for all three peptide substrates, suggesting a mechanism of BD association independent of the H3K4 modification state (**Fig. 3a**). CSPs were also seen in resonances for the PHD finger upon addition of all three substrates, but with unique bound state chemical shifts dependent on the H3K4 modification state. Specifically, H3tetra^ac^ and H3K9acK14acK18ac (H3tri^ac^) led to nearly identical bound state chemical shifts, contrasting with the unique signatures of H3K4me3tri^ac^ (**Fig. 3a**) Together this reveals that the PHD-BD associates with the acetylated H3 tail in a multivalent manner, employing both domains independent of H3K4 modification status, but forming a unique complex when H3K4 is trimethylated.

**Fig. 3.**
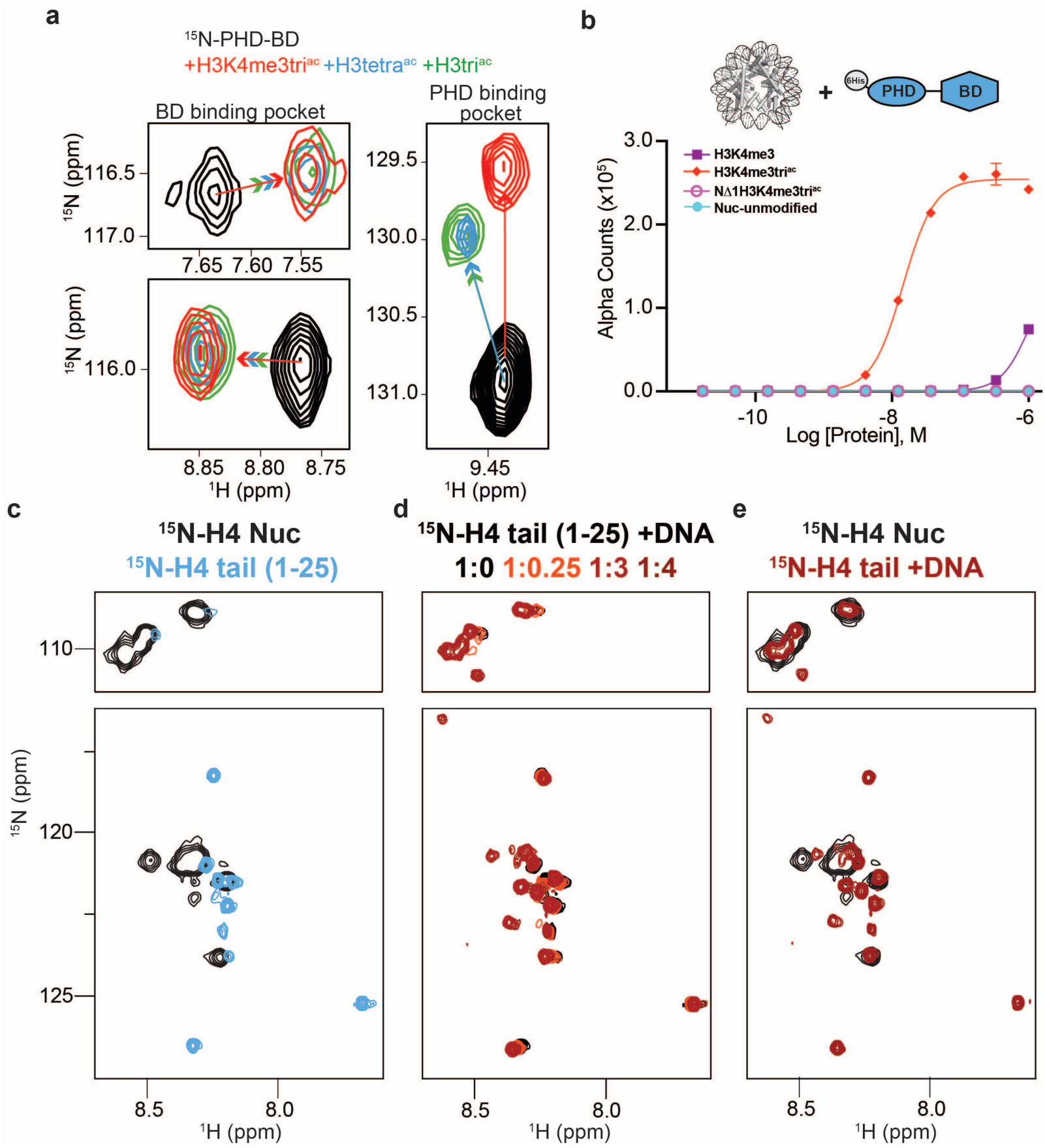
The preferred PTM binding pattern is dictated by Nuc conformation and multivalent reader binding potential. **a)** ^1^H,^15^N-HSQC overlays of ^15^N-PHD-BD apo (black) or in the presence of H3tri^ac^ (green), H3tetra^ac^ (blue), or H3K4me3tri^ac^ (red). Arrows denote trajectory of chemical shift perturbation and are colored by peptide. Shown are representative resonances for the BD (left) and PHD (right) binding pockets. **b)** Histone H3-A1 is essential for 6His-PHD-BD binding to Nucs (compare H3K4me3tri^ac^ to NΔ1H3K4me3tri^ac^). Alpha Counts from dCypher assays are plotted as a function of Query concentration to indicated Nuc Targets. **c)** Overlay ^1^H,^15^N-HSQC spectra of ^15^N-H4-Nuc (black) and ^15^N-H4-tail peptide (residues 1-25, blue). **d)** Overlay ^1^H,^15^N-HSQC spectra of ^15^N-H4-tail peptide upon titration of a 21bp double-stranded DNA. Molar ratios are denoted by color in legend. **e)** Overlay ^1^H,^15^N-HSQC spectra of ^15^N-H4-Nuc (black) and ^15^N-H4-tail peptide saturated with DNA (red).

The PHD/H3 binding interface includes pockets for H3 residues A1, R2, and K4me3^28^ (**Fig. 2a**), with K4me3 needed for a robust Nuc interaction in the isolated PHD context. From the above NMR data we hypothesized that in the tandem PHD-BD context the A1 and/or R2 interactions (even in the absence of K4me3) contribute to association with the acetylated H3 tail. To test this, we returned to dCypher and created Nucs with residue truncations at the H3 N-terminus (**Fig. 3b**), confirming their integrity by an antibody to H3K4me3 (**Extended Data Fig. 4d**). Upon H3 A1 deletion (NΔ1) PHD-BD had unmeasurable EC_50_^rel^ (**Fig. 3b**), indicating that recognition of the histone N-terminus is critical for stable Nuc binding (and consistent with the engagement mechanism for other PHD fingers^39^).

### DNA interactions occlude histone tail accessibility and lead to altered specificity

dCypher results showing a dramatically weaker (PHD-BD or PHD only) or undetectable (BD only) EC_50_^rel^ for Query binding to Nucs relative to peptides (*e*.*g*. **Extended Data Fig. 2**) are fully consistent with our previous NMR studies that revealed strong inhibition of PHD binding to H3K4me3 in the Nuc context^23^. There we demonstrated that H3 tail occlusion is due to interactions with nucleosomal DNA^23^, specifically proposing that the H3 tails adopt a high-affinity fuzzy complex driven largely by R/K residues^23,40^.

Interaction of the PHD-BD and BD alone with acetylated histone H4 was also abrogated in the Nuc context, despite robust binding to comparable peptides (**Fig. 1** and **Extended Data Figs. 1 - 2**). The H4 tail is K/R-rich, has decreased dynamics in the nucleosome vs. peptide context, and computational models suggest it may also form a fuzzy complex with DNA^41^. To further characterize H4 tail conformation in the Nuc environment, we utilized NMR spectroscopy to investigate a Nuc containing ^15^N-H4 (see **Methods**). Due to its large size (∼200 kDa) and resultant slow tumbling, it is expected that only very flexible regions (such as the tails) will be NMR observable using this isotope labeling scheme (**Fig. 1a**). Consistent with previous studies^41,42^ we observed resonances for only 15 of the 101 non-proline amino-acids of full length H4, corresponding to tail residues 1-15 (**Fig. 3c**). However, this represents only 15/20 possible resonances (assuming fast exchange on the NMR time-scale) for the H4 N-terminal tail (as classified by trypsin accessibility: **Fig. 1a**^43^). The severe line-broadening observed for residues 16-20 (also known as the H4 tail basic patch: **Extended Data Fig. 5**) indicated this region is likely stably associated with the Nuc core, in agreement with previous structural and biochemical studies^44^. However, the conformation of the observable first 15 residues are less clear.

We next generated the ^15^N-H4 tail in peptide form, allowing us to investigate conformational differences between a free tail and that in the Nuc context (as above). Overlay of the resulting spectra showed that every resonance has CSPs between peptide and Nuc (**Fig. 3c**), consistent with a different chemical environment for every residue. To determine if this is due to association with DNA, we collected sequential ^1^H,^15^N-HSQC spectra of the ^15^N-H4-tail peptide upon addition of unlabeled DNA (**Fig. 3d**). Notably, CSPs were seen for every resonance, revealing that the H4 tail bound DNA, and every tail residue is impacted. Overlay of the DNA-bound ^15^N-H4 tail spectrum with that for the ^15^N-H4-Nuc showed very similar chemical shifts, consistent with the entire H4 tail associating with nucleosomal DNA (**Fig. 3e**), which is in-line with previous cross-linking studies^45–47^. The differential linewidth of resonances indicates the H4 tail has two distinct dynamic regions: the first 15 residues exchanged quickly between multiple conformations on the DNA, consistent with a fuzzy complex^48,49^; while the basic patch exchanged much more slowly and/or between fewer states, leading to loss of signal. This is distinct from the H3 tail, where every residue experienced the fast dynamics of a fuzzy complex, and may be related to charge distribution and/or relative positioning with the NCP core.

The above data supports that, similar to the H3 tail, H4 tail conformation in the Nuc context occludes accessibility, and potentially explains the loss of BPTF BD and PHD-BD association with acetylated H4. To investigate if this conformation abrogated all interactions with the H4 tail we tested an alternate bromodomain Query (GST-BRD4-BD1; **Suppl. Tables 3: Resources A**) against peptides and Nucs (**Extended Data Fig. 6)**. BRD4-BD1 has previously been shown to bind acetylated H4 tail peptides^50^, and dCypher confirmed the strongest EC_50_^rel^ for H4tetra^ac^ over all peptides tested (**Extended Data Fig. 6a**,**c**). BRD4-BD1 also bound H4tetra^ac^ in the Nuc context, though with weaker affinity than the comparable peptide (EC_50_^rel^ [CI^95^] values of 7.4 nM [7.05-7.64] Nuc *vs*. 0.7 nM [0.64-0.73] peptide; **Extended Data Fig. 6b**,**c**). Thus, H4 tail accessibility is reader dependent (as also recently demonstrated for PHIP BD1-BD2^51^), and the ability to bind may rely on several factors including overall affinity or different engagement mechanisms. For instance, BRD4 BD1 (unlike BPTF BD) can associate with DNA^52^, and such competition may help disengage the H4 tail from the nucleosome core.

Together, this suggests that to enable association in the Nuc context a reader must be able to displace the modified histone tail from DNA. However, tail accessibility can be enhanced by disrupting the DNA interaction via modification of sidechain charge^21,23^, as where distal acetylation of the H3 tail improved BPTF-PHD engagement with H3K4me3 (**Extended Data Fig. 2a-d**). Notably, acetylation does not fully release the tail from DNA binding, as PHD still showed weaker association with the methylated/acetylated Nuc relative to peptide. This is consistent with previous studies indicating that acetylation weakens but does not fully disrupt histone tail DNA interactions^23,53^ .This may also explain why BPTF-BD alone was insufficient to establish binding with acetylated H3 and H4 tails.

### PHD-BD drives association of BPTF with the methylated and acetylated H3 tail *in vivo*

The above *in vitro* results indicate that BPTF PHD-BD associates with H3K4me3K14ac or H3K4me3K18ac (or H3K4me3K14acK18ac) in the Nuc context. To investigate if this preference is recapitulated *in vivo*, we performed CUT&RUN with antibodies to BPTF, H3K4me3, and H3K18ac (see **Methods** and **Suppl. Tables 3: Resources A**) in K562 cells. We observed extensive genomic co-localization of BPTF with each PTM, but the greatest degree of overlap when both are present (**Extended Data Fig. 7a-e**). As a bulk approach CUT&RUN is unable to confirm definitive co-enrichment of all elements, with one possible interpretation that our observations are due to distinct sub-populations. We thus designed a new approach (Reader CUT&RUN; see **Methods**) where GST-PHD-BD was complexed with an antibody to GST (α-GST) to create a CUT&RUN compatible reagent. We also developed DNA-barcoded PTM-defined Nucs (unmodified, H3K4me3, H3tetra^ac^ and H3K4me3tri^ac^; **Fig. 4a**) as a spike-in to monitor assay performance and GST-PHD-BD preference *in situ*. In these controlled studies GST-PHD-BD showed a dramatic preference for spike-in Nucs containing the combinatorial signature (H3K4me3tri^ac^) relative to each PTM alone (six-fold over H3K4me3, 41-fold over H3tetra^ac^; **Fig. 4b**), recapitulating the dCypher observations (*e*.*g*. **Fig. 1d**). The genomic enrichment of GST-PHD-BD further confirmed its combinatorial preference, with binding regions showing extensive overlap with those containing H3K4me3 and H3K18ac (**Fig. 4c-d**). Together with the spike-in results this is consistent with a synergistic association with both PTMs. Furthemore, the genomic enrichment of GST-PHD-BD was also highly correlated with that of endogenous BPTF (**Fig. 4c-d**), supporting that the tandem readers are sufficient to drive effective *in vivo* localization.

**Fig. 4.**
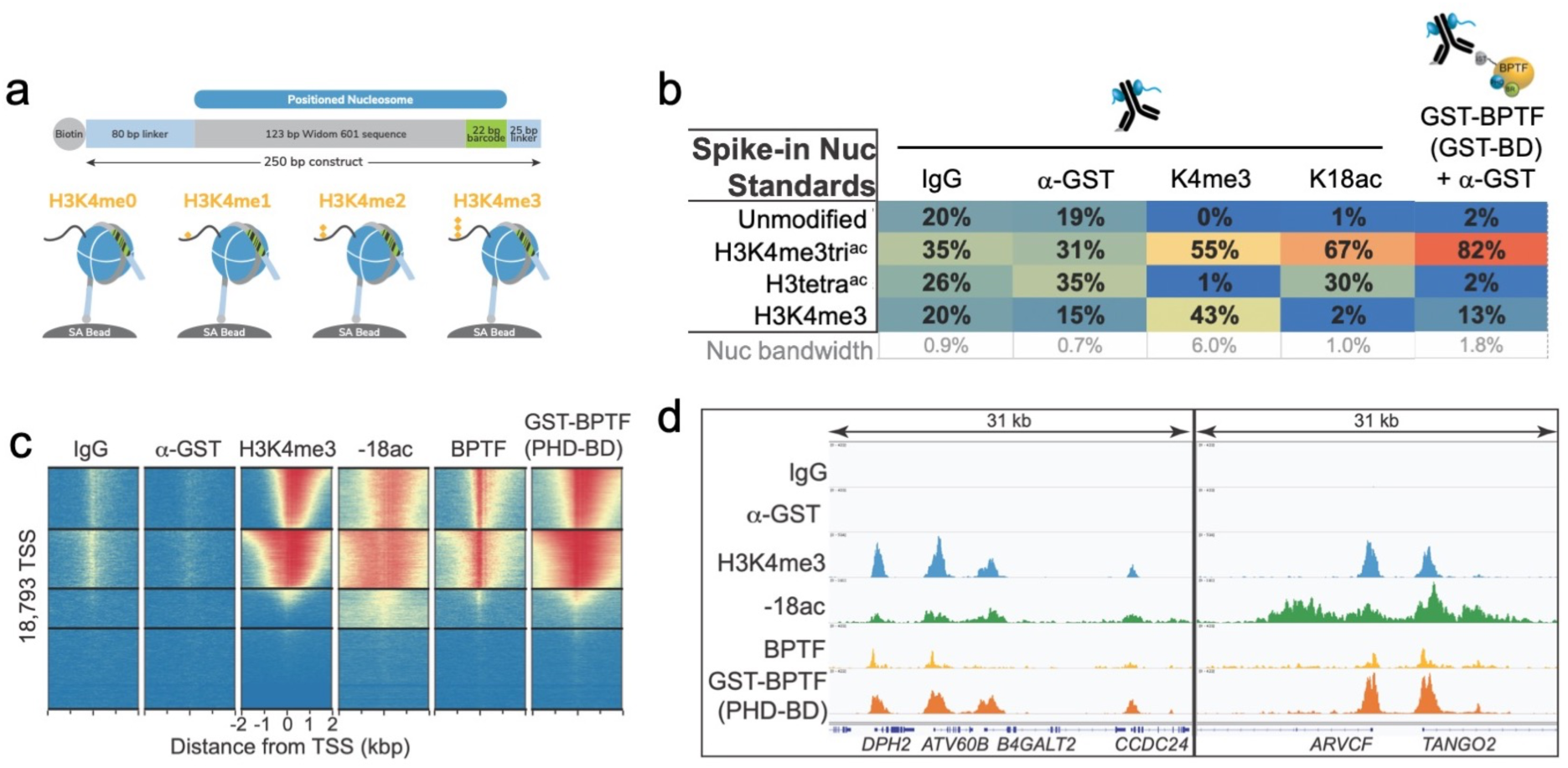
The *in vitro* combinatorial preference of BPTF PHD-BD is recapitulated *in vivo*. **a)** CUTANA Nuc spike-ins contain a 5’biotin for immobilization to magnetic beads and a DNA barcode to define PTM status / monitor release into the CUT&RUN eluate. A four-member panel was assembled to explore GST-PHD-BD binding (unmodified, H3K4me3, H3tetra^ac^, H3K4me3tri^ac^; on 80-<u>N</u>-25 DNA containing a central 147bp 601 <u>N</u>ucleosome positioning sequence with embedded 22bp DNA barcode). **b)** GST.PHD-BD shows strong preference for spike-in Nucs containing H3K4me3tri^ac^. Table shows relative release of spike-ins (percent barcoded Nuc / total barcode reads) in Reader CUT&RUN (**Methods**). Antibodies are noted by column; GST-BPTF (PHD-BD) is detected by α-GST to facilitate pAG-MNase recruitment. ‘Nuc bandwidth’ is the percentage of total sequence reads taken up by Nuc spike-in standards. **c)** Heatmap of CUT&RUN signal aligned to the transcription start site (TSS, +/-2kb) of 18,793 genes in K562 cells. Rows were k-means clustered into four groups (boxed) using ChAsE chromatin analysis tool^92^. High and low signal (red and blue respectively) are ranked by / linked to H3K4me3 (top to bottom). **d)** CUT&RUN RPKM normalized tracks at representative loci using Integrative Genomics Viewer (IGV, Broad Institute). Note the co-localization of BTPF (endogenous) or GST-PHD-BD (exogenous) with H3K4me3 and H3K18ac; that H3K18ac alone is insufficient to recruit BTPF or GST-PHD-BD; and that GST-PHD-BD shows robust recruitment at some locations where BPTF is absent (*e*.*g. B4GALT2* promoter; see **Discussion**).

## DISCUSSION

Taken together, our data indicates that nucleosome context strongly influences reader domain engagement with histone PTMs. Previous studies have noted reduced reader affinity on Nucs relative to histone tail peptides^23–26^, but here we show that the PTM(s) bound may also be restricted (*e*.*g*. loss of Nuc H4acetyl binding by the BPTF BD and PHD-BD tandem; **Fig. 1c-e** and **Extended Data Fig. 2a-b, f-g**), or the preferred modification pattern may be altered (*e*.*g*. the PHD preference for Nucs with H3K4me3 and additional tail acetylation; **Fig. 2b,e**). We propose this is due (at least in part) to association of the histone tails with nucleosomal DNA. This conformation limits accessibility and leads to competition for the tails between the DNA and reader domain(s). As a result, histone PTMs may play multiple roles; weakening the DNA association to increase access for reader domains, providing a platform for reader domain binding, or both.

As a result of occluded tail conformation in the Nuc context, the multivalent binding of tandem domains is not simply defined by raw potential (*i*.*e*. the sum of individually preferred PTMs), but also binding opportunity. For BPTF PHD-BD, this manifests as a nucleosomal restriction on H4ac tail binding, leaving selectivity for H3ac. We observe multiple ways to combine multivalent contacts across domains, and thus support productive engagement. In the case of the BPTF PHD-BD tandem, the PHD can associate with H3 A1, R2 and K4me3 (**Fig. 3a-b**), while the BD can bind K14ac and K18ac (**Extended Data Fig. 1f-g**). Notably, when the H3 tail is only acetylated (as in the H3tetra^ac^ Nuc) the resulting weakening of the tail / DNA interaction combined with BD binding to Kac and PHD finger binding to A1 and R2 together support weak Nuc engagement. Alternatively, for H3K4me3 absent any acetylation, PHD contacts with A1, R2 and K4me3 also support weak Nuc engagement. Finally, strong binding occurs when H3K4me3 and H3K14ac or H3K18ac are present, promoting tail displacement and allowing both the PHD and BD to most effectively engage. The preference of BPTF PHD-BD for H3K4me3 with H3K14ac or H3K18ac over H3K9ac may be due to the *in cis* proximity of K9 to H3K4me3, restricting BD binding when the PHD finger is engaged. Thus, within a tail displacement model, tandem domains can accommodate multiple distinct PTM signatures to engage modified Nucs. Notably, and as seen here, these may have varying strengths of interaction which in turn may mediate an array of responses within the chromatin landscape, including differences in CAP retention at specific sites, or stabilization at an intermediate modified state.

When moving from the peptide to Nuc context we (and others) consistently observe individual reader domains to show reduced affinity and restricted specificity^23–26,35,51^. An exception to this is readers with intrinsic DNA binding ability, such as the PWWPs. These form multivalent interactions with DNA and histone tails (so peptide studies are often uninformative)^54– 60^, but may also act to directly compete for the DNA, thus promoting PTM accessibility on their target tail^61^. Indeed, several mechanisms for modulating histone tail conformation can be imagined^40^. Beyond *in cis* modification of the target histone tail (as in this study), modification of an adjacent tail may alter the dynamics of the target, such *trans*-tail crosstalk being recently reported for H3 and H4^62^. Adjacent DNA binding domains within the same protein or complex may also play a role in displacing the target tail from DNA. Alternatively, histone tail accessibility can be modulated by changes to the canonical nucleosome composition, such as hexasomes depleted of one H2A-H2B dimer^63^.

In reader-CUT&RUN GST-PHD-BD recapitulated the dCypher preference for spike-in Nucs containing the combinatorial target (H3K4me3tri^ac^) over each PTM class alone (**Fig. 4b**). Furthermore, GST-PHD-BD localization across the genome was highly correlated with regions that also contain H3K4me3, H3K18ac and endogenous BPTF (**Figs. 4c-d**). Together, this suggests that the combinatorial readout of these PTMs is indeed a discerning factor in the genomic localization of BPTF: the activity of both domains is clearly important to achieve robust interaction, and thus at minimum critical to achieve proper kinetics on chromatin.

In an extended analysis of our genomics data, we considered that full-length BPTF (endogenous) could harbor additional regulatory potential over exogenous BPTF PHD-BD. In this regard while the dominant signature was where H3K4me3 / H3K18ac co-localized with both endogenous and exogenous (**Fig. 4c-d**), we observed numerous locations where the PTM combinatorial overlapped only with exogenous (as at *B4GALT2* in **Fig. 4d**), while the contrasting pattern (PTMs only overlapped with endogenous) was a much rarer species. This may be due to the relative level of exogenous to endogenous protein (or the target PTMs), where one might expect a higher abundance exogenous to extend to locations of lower PTM density. However peak structure comparison does not appear to support this explanation, as sites retaining exogenous but lacking endogenous are not the weakest H3K4me3 / H3K18ac locations. We speculate the more interesting possibility: endogenous BPTF is subject to regulation that further refines its chromatin localization beyond simple availability of H3K4me3 / H3K18ac for its C-terminal PHD-BD. Indeed, there are increasing examples of auto-regulatory elements within CAPs that modulate their activity^64–78^, suggesting that a histone code is more than the simple availability of potentially redundant positive signals.

It is becoming increasingly clear that we should interrogate the binding of readers to histone PTMs with more physiological entities: moving away from minimal-domain queries and histone peptide targets to full length CAPs (or higher order complexes) and Nucs, and thus accommodate the regulatory potential on each side. Doubtless a more thorough mechanistic understanding will reveal novel approaches to target these interactors with therapeutic intent.

## Supporting information

Supplementary Discussion and Figures

## AUTHOR CONTRIBUTIONS

MRM, IKP, AV and NWH designed, performed and analyzed *dCypher* studies. MJM, RW, SAH and MAC created / validated PTM-defined histones, octamers and nucleosomes. JMB created versaNucs with peptides validated by SAH. KN, EM and BJV performed CUT&RUN studies. HAF performed NMR and MS analyses under supervision by CM. ZWS and MCK supervised research at *EpiCypher*. MCK and CM co-wrote the manuscript with contributions / support from all authors.

## ACKNOWLEDGEMENTS

HAF was supported by an NIH T32 fellowship (2T32GM008365-26A1) through the Center for Biocatalysis and Bioprocessing. This project was supported by The Holden Comprehensive Cancer Center at The University of Iowa and its National Cancer Institute Award (P30CA086862). Work in the Musselman lab was funded by grants from the National Science Foundation (CAREER-1452411) and the National Institutes of Health (NIH; R35GM128705). *EpiCypher* is supported by NIH grants R44GM117683, R44CA214076, R44GM116584 and R44DE029633. We would like to thank Vic Parcell and the High-Resolution Mass Spectrometry Facility (Office of the Vice-President for Research and Economic Development at the University of Iowa) for technical support.

## METHODS

### BPTF protein constructs and preparation

Human BPTF (<u>Uniprot Q12830</u>) PHD finger-bromodomain (PHD-BD) and PHD finger were cloned into pGEX6p with an N-terminal Glutathione S-Transferase (GST) tag and a PreScission Protease cleavage site (**Suppl. Tables 3: Resources A** and **Extended Data Fig. 9**). BPTF BD with an N-terminal 6xHistidine (6His) tag and Tobacco Etch Virus (TEV) Protease cleavage site was from *Addgene* (<u>plasmid 39111</u>). This was modified using the Q5 Site-directed mutagenesis kit (*New England Biolabs* [*NEB*]) for domain addition / removal or single amino acid substitutions. All constructs were expressed in *E*.*coli* BL21 (DE3) (*ThermoFisher Scientific* or *NEB*). Cells were grown to OD_600_ ∼1.0 and induced with 0.8 mM IPTG at 18°C for ∼16 hr in LB (or M9 minimal media for NMR). M9 media was supplemented with vitamin (*Centrum* Adult), 1 g/L ^15^NH_4_Cl, and 5 g/L D-glucose. For constructs containing the BPTF PHD finger all growth media and buffers were supplemented with 100 µM ZnCl_2_. For purification of BPTF recombinants cells were lysed by sonication, and lysates incubated with either glutathione agarose (*Thermofisher Scientific*) or Ni-NTA resin (*Thermofisher Scientific*) to respectively enrich for GST- and 6His-tagged proteins. Fusion proteins were eluted with reduced L-glutathione or imidazole as appropriate. For NMR, samples were cleaved from the GST tag using PreScission Protease. All BPTF proteins were then further purified using anion exchange (Source 15Q, *GE Healthcare Life Sciences*) and size exchange chromatography (Superdex 75, *GE Healthcare Life Sciences*). Protein concentrations were determined by UV-Vis spectroscopy.

### Histone preparation and nucleosome core particle reconstitution for NMR

Unmodified human histones H2A, H2B, and H3 (**Resource Table A**) were expressed in *E*.*coli* Rosetta 2 (DE3) *pLysS* or BL21 (DE3) in LB media. Cells were grown to OD_600_ ∼0.4 and induced with 0.4 mM IPTG at 37°C for either 3 hr (for H3) or 4 hr (for H2A and H2B). ^15^N-labelled histone H4 (**Suppl. Tables 3: Resources A**) was expressed in Rosetta 2 (DE3) *pLysS* cells from a pET3a vector in M9 minimal media supplemented with vitamin, 1 g/L ^15^NH_4_Cl, and 5 g/L D-glucose. Cells were induced at OD_600_∼0.4 with 0.2 mM IPTG at 37°C for 3 hr. Histones were purified from inclusion bodies as previously^79^ and purified by ion exchange. Mass spectrometry with positive electrospray ionization (Waters Q-Tof Premier instrument) was used to validate the histones and ensure no carbamylation occurred during purification (**Extended Data Fig. 8**). Samples were diluted 1:2 or 1:4 in water/acetonitrile (1:1) with 0.1% (v/v) formic acid. The acquisition and deconvolution software used during data collection and analysis were MassLynx and MaxEnt, respectively.

Histone octamers were prepared as described^79^. In brief, equimolar ratios of purified histones were combined in 20 mM Tris pH 7.5, 6M Guanidine HCl, 10 mM DTT and subsequently dialyzed into 20 mM Tris pH 7.5, 2M KCl, 1 mM EDTA, 5 mM β-mercaptoethanol (β-ME). Octamers were purified via size chromatography over a Sephacryl S-200 column (*GE Healthcare Life Sciences*).

The 147 bp Widom 601 nucleosome positioning sequence (NPS)^80^ was amplified in *E*.*coli* using a plasmid containing 32 repeats (**Suppl. Tables 3: Resources A**). DNA was purified by alkaline lysis^79^, the 147bp 601 NPS excised from the plasmid with *EcoRV*, polyethylene glycol precipitated, and further purified over a source 15Q column (*GE Healthcare Life Sciences*).

Reconstitution of Nucleosome core particles (NCPs; Nucs) with the 147 bp DNA was accomplished by desalting^79^. In brief, octamer and DNA were combined in equimolar amounts in 2M KCl and desalted to 150 mM KCl using a linear gradient over ∼48 hr. Nucs were heat-shocked at 37°C for 30 min for optimal positioning and purified using a 10-40% sucrose gradient. Nuc formation was confirmed by sucrose gradient profile and native polyacrylamide gel electrophoresis (see **Extended Data Fig. 8**). Nuc concentrations were determined by UV-vis spectroscopy (after diluting in 2M KCl to disassemble NCPs) using the absorbance from 601 DNA (calculated ε_260_= 2,312,300.9 M^-1^cm^-1^).

### H4 tail peptide purification for NMR

The histone H4 tail (residues 1-25 followed by a C-terminal tyrosine for quantification) was expressed from pGEX6p as a fusion protein with an N-terminal GST tag followed by a PreScission Protease cleavage site (**Suppl. Tables 3: Resources A**). This was overexpressed in *E*.*coli* BL21 (DE3) (*NEB*) grown in M9 minimal media supplemented with vitamin (Centrum daily multivitamin), 1 g/L ^15^NH_4_Cl, and 5 g/L D-glucose. Cells were grown to an OD_600_∼1.0 and induced with 0.5 mM IPTG at 37°C for 4 hr. The ^15^N-GST-H4 peptide fusion was purified on glutathione agarose resin (*Thermofisher Scientific*), cleaved with PreScission Protease (16 hr incubation at 4°C), and products resolved by size exclusion chromatography (Superdex 75 10/300; *GE Healthcare Life Sciences*). Peptide identity was validated by mass spectrometry with positive electrospray ionization (on a Waters Q-Tof Premier). Samples were diluted 1:2 or 1:4 in water/acetonitrile (1:1) with 0.1% formic acid. The acquisition and deconvolution software used during data collection and analysis were MassLynx and MaxEnt, respectively. ^15^N-H4 (1-25) peptide concentration was determined by UV-vis spectroscopy using the non-native C-terminal tyrosine.

### DNA preparation for NMR

Oligonucleotides (5’-CTCAATTGGTCGTAGACAGCT-3’ and the complement 5’-AGCTGTCTACGAACCAATTGAG-3’) for DNA titration NMR were from *Integrated DNA Technologies* (*IDT*). These were annealed at 50 µM by heating to 94°C followed by gradual cooling to room temperature (in 10 mM Tris-HCl pH 7.5, 50 mM NaCl, 1 mM EDTA). Duplex DNA was purified by ion exchange chromatography on a source 15Q column (*GE Healthcare Life Sciences*) and analyzed by 1% agarose gel. DNA was precipitated in ethanol, resuspended in ddH_2_O, and concentration determined by UV-vis spectroscopy and the predicted extinction coefficient (ε_260_= 333,804.5 M^-1^ cm^-1^).

### NMR spectroscopy

^1^H-^15^N heteronuclear single quantum coherence (HSQC) spectra were collected on 30 µM ^15^N-H4 tail peptide and 80.5 µM nucleosome samples in 20 mM MOPS pH 7.2, 150 mM KCl, 1 mM DTT, 1 mM EDTA, and 10% D_2_O. Data was collected at 25°C on an 800 MHz Bruker spectrometer equipped with a cryoprobe. Titration of the 21 bp dsDNA into ^15^N-H4 tail peptide was performed through the collection of sequential ^1^H-^15^N HSQC spectra on the _15_N-H4 tail in the apo state and with increasing DNA concentrations (spectra collected at [peptide:DNA] molar ratios of 1:0, 1:0.1, 1:0.25, 1:0.5, 1:1, 1:2, and 1:3).

Sequential ^1^H-^15^N HSQC spectra of 25 µM ^15^N-BD and ^15^N-BD (N3007A) were collected with increasing concentrations of H4K16ac tail peptide (**Suppl. Tables 3: Resources D**) in 50 mM potassium phosphate pH 7.2, 50 mM KCl, 1 mM DTT, 1 mM EDTA, and 10% D_2_O at 25°C on an 800MHz Bruker spectrometer equipped with a cryogenic probe. Concentration of the stock H4K16ac peptide was analyzed by Pierce Quantitative Fluorometric Peptide Assay (*Thermofisher Scientific*). Spectra were collected with [^15^N-BD: H4K16ac peptide] at ratios 1:0, 1:0.5, 1:1, 1:2.5, 1:5, 1:10, 1:20, 1:30, 1:50, 1:70 and [^15^N-BD (N3007A): H4K16ac peptide] at ratios 1:0, 1:5, 1:20, 1:40.

Sequential ^1^H-^15^N HSQC spectra of 50 µM ^15^N-PHD-BD were collected with increasing concentrations of histone tail peptides (H3K4me3tri^ac^, H3tetra^ac^ or H3tri^ac^) were collected in 50 mM potassium phosphate pH 7.2, 50 mM KCl, 1mM DTT, 25 µM ZnCl_2_, and 10 % D_2_O. Experiments were collected at 25°C on an 800 MHz Bruker spectrometer equipped with a cryogenic probe. Data was collected for [^15^N-PHD-BD: H3 tail peptide] at ratios 1:0, 1:0.1, 1:0.5, 1:1, 1:2, 1:4, and 1:8. All NMR data was processed using NMRPipe^81^ and analyzed using CcpNmr Analysis^82^.

### Histone peptides for *dCypher*

All histone peptides for *dCypher* (**Suppl. Tables 3: Resources B**) were synthesized with a terminal Biotin (location as indicated) and identity confirmed by mass spectrometry.

### Semi-synthetic nucleosomes with defined post-translational modifications (PTMs)

PTM-defined histones, octamers and nucleosomes [dNucs™ or versaNucs^®^] for *dCypher* were synthesized / purified / assembled as previously^83,84^ but without DNA barcoding (see **Suppl. Discussion**; **Suppl. Tables 3: Resources C - D**; and **Extended Data Fig. 9**).

### *dCypher* binding assays

*dCypher* binding assays to PTM-defined peptides / Nucs were performed as previously (see **Suppl. Discussion**)^59,85^. In brief 5 μl of GST-or 6HIS-tagged reader domain (Query: specific identity / concentration as indicated) was incubated with 5 μl of biotinylated peptide (100 nM final) / Nuc (10 nM final) (Target: specific identity as indicated) for 30 min at room temperature in the appropriate assay buffer ([Peptide: 50mM Tris pH 7.5, 50mM NaCl, 0.01% Tween-20, 0.01% BSA, 0.0004% Poly-L Lysine, 1mM TCEP]; [Nuc: 20mM HEPES pH 7.5, 250mM NaCl, 0.01% BSA, 0.01% NP-40, 1mM DTT]) in a 384-well plate. For GST-tagged proteins a 10 μl mix of 2.5 μg/ml glutathione (*PerkinElmer*) and 5 μg/ml streptavidin donor beads (*PerkinElmer*) was prepared in peptide or Nuc bead buffer ([Peptide: as assay buffer]; [Nucs: as assay buffer minus DTT]) and added to each well. For 6HIS-tagged proteins a 10 μl mix of 2.5 μg/ml Ni-NTA acceptor beads (*PerkinElmer*) and 10 μg/ml streptavidin donor beads was used. The plate was incubated at room temperature in subdued lighting for 60 min and the Alpha signal measured on a *PerkinElmer 2104 EnVision* (680-nm laser excitation, 570-nm emission filter ± 50-nm bandwidth). Each binding interaction was performed in duplicate^86^.

Binding curves [Query : Target] were generated using a non-linear 4PL curve fit in Prism 8 (*GraphPad*), with EC_50_^rel^ values and 95% confidence intervals (CI^95^) computed and converted from log(X) using antilog 10^X (see **Suppl. Discussion** and **Suppl. Tables 1 & 2**). Where necessary, values beyond the Alpha hook point (indicating bead saturation / competition with unbound Query)^86^ were excluded and top signal constrained to average max signal for Target (in cases where signal never reached plateau, those were constrained to the average max signal within the assay). For statistical analysis, unpaired two-tailed t-tests were performed in Prism using Log(EC_50_^rel^) and standard error values / differences considered statistically significantly when P< 0.05 (see **Suppl. Tables 1 & 2**).

### CUTANA CUT&RUN, Illumina sequencing, and data analysis

CUT&RUN was performed with native or fixed (H3K18ac only; see below) K562 cells using CUTANA™ protocol **v1.5.1**^87^ which is an optimized version of that previously described^88^. For each native CUT&RUN reaction, 500K permeabilized cells were immobilized onto Concanavalin-A beads (Con-A; *EpiCypher* #21-1401) and incubated overnight (4° C with gentle rocking) with 0.5 µg of antibody (IgG, H3K4me3, H3K18ac and BPTF [**Suppl. Tables 3: Resources E**; all PTM antibodies validated to SNAP-ChIP Nuc standards as previously^83^]). pAG-MNase (*EpiCypher* #15-1016) was added / activated and CUT&RUN enriched DNA purified using the *Monarch DNA Cleanup* kit (*NEB* #T1030S). 10 ng DNA was used to prepare sequencing libraries with the *Ultra II DNA Library Prep* kit (*NEB* #E7645S).

Some labile PTMs benefit from a light fixation step (not shown), so minor protocol modifications were made for H3K18ac. 500K cells were crosslinked with 0.1% formaldehyde for 1 minute at room temperature, and then quenched with 125 mM glycine. To help the cellular ingress of antibody / egress of cleaved chromatin fragments the Wash, Antibody, and Digitonin buffers were supplemented with 1% Triton X-100 and 0.05% SDS. To reverse crosslinks prior to DNA column cleanup, CUT&RUN eluate was incubated overnight at 55° C with 0.8 µl 10% SDS and 20 µg Proteinase K (*Ambion* #AM2546).

Libraries were sequenced on the Illumina platform, obtaining ∼ 4 million paired-end reads on average (**Suppl. Tables 3: Resources E**). Paired-end fastq files were aligned to the hg19 reference genome using the Bowtie2 algorithm^89^. Only uniquely aligned reads were retained, and blacklist regions^90^ filtered out prior to subsequent analyses. Peaks were called using SEACR (<u>S</u>parse <u>E</u>nrichment <u>A</u>nalysis of <u>C</u>UT&<u>R</u>UN)^91^. All sequencing data has been deposited in the NCBU Gene Expression Omnibus (GEO) with accession number <u>GSE150617</u>.

### Reader CUT&RUN

Reader CUT&RUN (*i*.*e*. GST-PHD-BD) was performed as above for CUTANA CUT&RUN with the following modifications. 500K native K562 cells were used for each reaction and all buffers were supplemented with 1 µM TSA (Trichstatin A, *Sigma* #T8552) to protect potentially labile acetyl-PTMs (*e*.*g*. H3K18ac).

A biotinylated CUTANA Nuc mini-panel (unmodified, H3K4me3, H3tetra^ac^, H3K4me3tri^ac^; each on 80-<u>N</u>-25 DNA containing a central 147bp Widom 601 <u>N</u>ucleosome positioning sequence with embedded 22bp DNA barcode: **Fig. 4a-b**) was synthesized, individually coupled to magnetic streptavidin beads (*NEB #S1421S*) at saturation, and spiked into each CUT&RUN reaction (final concentration 0.8 nM) with Con-A immobilized cells just prior to antibody addition. Each member of the Nuc panel was DNA barcoded to define PTM status / monitor comparative release into the CUT&RUN eluate (to be quantified after sequencing). After Nuc spike-in, GST-PHD-BD or GST (**Suppl. Tables 3: Resources B**) and IgG (**Suppl. Tables 3: Resources E**) were added to the same sample (final concentration 70nM each), and incubated overnight at 4° C. Samples were washed twice, and then incubated with 0.5 µg anti-GST (**Suppl. Tables 3: Resources E**) at room temperature for 30 min. The remainder of the assay was performed using the standard CUT&RUN protocol and sequenced as above. All sequencing data has been deposited in GEO with accession number <u>GSE150617</u>.

#### ABBREVIATIONS

BD: Bromodomain
BPTF: Bromodomain
PHD: Finger Transcription Factor
CAP: Chromatin Associated Protein
CSP: Chemical shift perturbation
Nuc: Nucleosome
PTM: Post-Translational Modification

