## Supplementary Discussion and Figures for "Nucleosome conformation dictates the histone code"

### CONTENTS OF THIS DOCUMENT


| | |
|---|---|
| Supplementary Discussion | p3 - 4 |
| Supplementary Figures | p5 - 13 |
| Supplementary References | p14 - 15 |


---

### ADDITIONAL SUPPLEMENTARY DOCUMENTS

**Suppl. Tables 1 & 2_Marunde et al.xlsx**

**Tab 1: Supplementary Table 1**. All $EC_{50}^{rel}$ in study [Figure by Figure + Extras]

**Tab 2: Supplementary Table 2**. 6HIS vs GST tagged BPTF queries, with trend similarities and accessibility / co-operative engagement. This info is referenced throughout study and assembled here for quick comparison.

**Tab 3:** T-test results (for specific comparisons in text).

**Tabs 4 - 18:** Raw data for figures. Associated with $EC_{50}^{rel}$.

**Suppl. Tables 3_Marunde et al.xlsx**

**Resources A:** Plasmids + Proteins

**Resources B:** dCypher Peptides (287x)

**Resources C:** dCypher dNucs (59x)

**Resources D:** Peptides (versa, 18x) & NMR

**Resources E:** CUT&RUN antibodies (and Sequence stats)

---

### SUPPLEMENTARY DISCUSSION

*The dCypher approach and calculation of $EC_{50}^{rel}$*

The *dCypher*® approach (*dCypher* for brevity) was developed on the chemiluminescent bead-based, no-wash Alpha platform (*PerkinElmer*) for the high-throughput profiling of Chromatin-Associated Protein (CAP) binding to PTM-defined histone peptides and semi-synthetic nucleosomes (Nucs)[1–7]. In brief, biotinylated peptides or Nucs (the potential **Targets**) were individually coupled to streptavidin-coated "Donor" beads, while epitope-tagged proteins (**Queries**) were bound to anti-tag "Acceptor" beads. After mixing potential reactants in a 384-well format, the Donor beads were excited at 680 nm, releasing singlet oxygen that caused emission (520–620 nm) in proximal (within 200 nm) Acceptor beads; this luminescent signal is directly correlated to interaction / binding affinity. A complete description of the *dCypher* approach is available[5,6].

To rank [Query : Target] binding we used a four-parameter logistical (4PL) model and reported the resulting data fit as relative $EC_{50}$s ($EC_{50}^{rel}$)[8] and 95% confidence intervals ($CI^{95}$) (see **Suppl. Tables 1 & 2** for all from this study). These values are defined as the concentration of Query (*e.g.* GST-PHD-BD) required to provoke a half-maximal response to Target along a representative dose-response curve[9]. Notably, we report as $EC_{50}^{rel}$ because a stable maximal response (100% ±5%) control is not included during data generation: as such we cannot ensure saturation. Although these values can be directly compared across Queries to understand relative binding (as they are within this study), they are not treated as an equilibrium dissociation constant ($K_d$). Very specific parameters must be met within the set-up of an Alpha assay to define a binding interaction $K_d$ : namely the generation of saturation curves, or a competition assay to identify the Query concentration at least 5x below bead binding saturation using 10x Target[10].

*Comparison of affinity tags [GST vs. 6His]*

For *dCypher* assays in this study the BPTF PHD-BD and PHD Queries were N-terminally tagged with either GST- or 6His-, while the BD Query was only available as 6His- (due to expression / purification difficulties).

A comparison of both epitope-tagged forms of PHD-BD revealed the resulting $EC_{50}^{rel}$ data for most homologous Targets to rank order identically (**Suppl. Tables 1 & 2**). One interesting exception was for PHD-BD binding to H3K4me1, me2, and me3 peptides. In this case, GST-PHD-BD showed similar $EC_{50}^{rel}$ for each H3K4 methyl state, while 6His-PHD-BD displayed a moderate preference for H3K4me3. Furthermore, the $EC_{50}^{rel}$ values of GST-tagged Queries were reduced compared to their 6His-tagged counterparts, indicating tighter binding. These results may be due to the use of epitope-specific beads (*i.e.* glutathione *vs.* nickel chelate acceptor beads; with the former being potentially more sensitive), and/or the dimerization of GST[11], which would be expected to enhance [Query : Target] binding via an effective local increase in Query concentration. Given the above, we note the importance to only compare $EC_{50}^{rel}$ values between similarly tagged Queries.

*Semi-synthetic Nucleosomes*

All Nucs in this study were from the *EpiCypher* dNuc™ or versaNuc® portfolios. For dNucs, PTM-defined histones were mixed (at mg scale) to a defined stoichiometry and dialyzed / purified to octamers, which were subsequently assembled on 147bp 5' biotinylated 601 DNA[12]. The resulting products (e.g. H3K4me3; *#16-0316*) contained full-length 'scarless' histones and minimal free-DNA (<5%). For versaNucs, histone H3 tail peptides (aa1-31 (A29L) with a PTM (or mutation) of interest were individually ligated to a H3 tailless nuc precursor (H3.1NΔ32 assembled on 147bp 5' biotinylated 601 DNA; *#16-0016*). The resulting Nucs (assembled at 50-100μg scale) contained minimal free DNA (<5%), undetectable levels of peptide precursor, and ≥90% full-length H3.1 with the PTM(s)/mutations of interest (*e.g.* **Extended Data Fig. 9**)[13]**.** In general, we observed no discernible difference in dNuc and versaNuc behavior (not shown), and so they are used interchangeably in this study (while always including both forms if available; **Suppl. Tables 3: Resources C - D**). However, versaNucs are not recommended for studies that encroach on the A29L position (as present in the final product): *e.g.* for modifiers or binders to H3R26, K27 or S28.

### SUPPLEMENTARY FIGURES

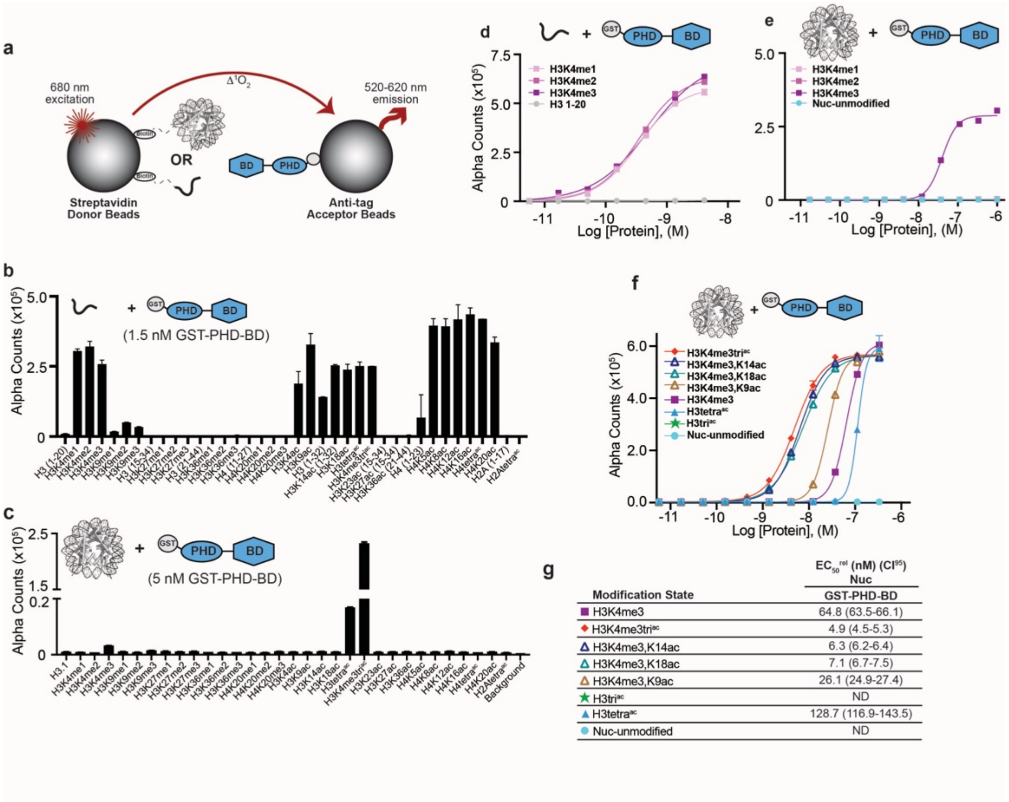


| Modification State | EC50rel (nM) (CI95) Nuc GST-PHD-BD |
|---|---|
| H3K4me3 | 64.8 (63.5-66.1) |
| H3K4me3tri^ac | 4.9 (4.5-5.3) |
| H3K4me3,K14ac | 6.3 (6.2-6.4) |
| H3K4me3,K18ac | 7.1 (6.7-7.5) |
| H3K4me3,K9ac | 26.1 (24.9-27.4) |
| H3tri^ac | ND |
| H3tetra^ac | 128.7 (116.9-143.5) |
| Nuc-unmodified | ND |

**Extended Data Fig. 1. Binding preference of BPTF GST-PHD-BD for peptide vs. Nuc substrates. a)** dCypher® is a no-wash bead-based proximity assay on the Alpha platform to interrogate chromatin readers (see **Methods** and **Suppl. Discussion**). PTM-defined biotinylated peptides or Nucs (the potential Targets) are immobilized on streptavidin donor beads and tagged reader domain(s) (the Query) bound to anti-tag Acceptor beads. Donor beads are excited at 680nm causing the release of singlet oxygen and quantitative fluorescence emission (520-620nm) from proximal acceptor beads (*i.e.* bridged by a [Query: Target] interaction). **b,c)** GST-PHD-BD selectivity (Query concentration as noted; chosen from dCypher curves in **Fig. 1d,e** [$EC_{80}^{rel}$ for H3K4me3 peptide; $EC_{50}^{rel}$ for H3K4me3tri$^{ac}$ Nuc]) for a panel of histone peptides (~100 nM final; **b**) and Nucs (10 nM final; **c**). **d,e)** GST-PHD-BD binding to histone H3 lysine 4 (H3K4) methyl states (me0 / me1 / me2 / me3) on peptides (**d**) or Nuc (**e**). Alpha Counts are plotted as a function of Query concentration. **f)** Alpha counts plotted as a function of GST-PHD-BD concentration from dCypher assays with Nucs. **g)** $EC_{50}^{rel}$ values with 95% confidence intervals ($CI^{95}$) (see **Methods** and **Suppl. Discussion** for calculation of same) from dCypher curves in (**f**) with Targets color coded as per legend (see also **Suppl. Tables 1 & 2**). ND, Not Detected.

a

| Modification State | $EC_{50}^{rel}$ (nM) ($CI^{95}$) Peptide 6His-PHD | 6His-BD |
|---|---|---|
| ■ H3K4me3 | 42.0 (39.1-44.8) | - |
| ◆ H3K4me3tri$^{ac}$ | 100.0 (96.2-104.2) | >1000 |
| ▲ H3tetra$^{ac}$ | ND | >1000 |
| ★ H3tri$^{ac}$ | - | - |
| ● H4tetra$^{ac}$ | - | 224.4 (215.8-233.3) |
| ● H3 1-20 | >1000 | ND |
| ● H4 1-23 | - | ND |

b

| Modification State | $EC_{50}^{rel}$ (nM) ($CI^{95}$) Nuc 6His-PHD | 6His-BD | GST-PHD |
|---|---|---|---|
| ■ H3K4me3 | >1000 | ND | 345.8 (344.3-347.5) |
| ◆ H3K4me3tri$^{ac}$ | 325.3 (315.5-335.0) | ND | 38.7 (38.3-39.1) |
| △ H3K4me3,K14ac | - | - | 98.0 (92.0-104.2) |
| △ H3K4me3,K18ac | - | - | 114.7 (107.4-122.7) |
| △ H3K4me3,K9ac | - | - | 101.1 (95.7-106.9) |
| ▲ H3tetra$^{ac}$ | ND | ND | - |
| ★ H3tri$^{ac}$ | - | ND | ND |
| ● H4tetra$^{ac}$ | - | ND | - |
| ● Nuc-unmodified | ND | ND | ND |

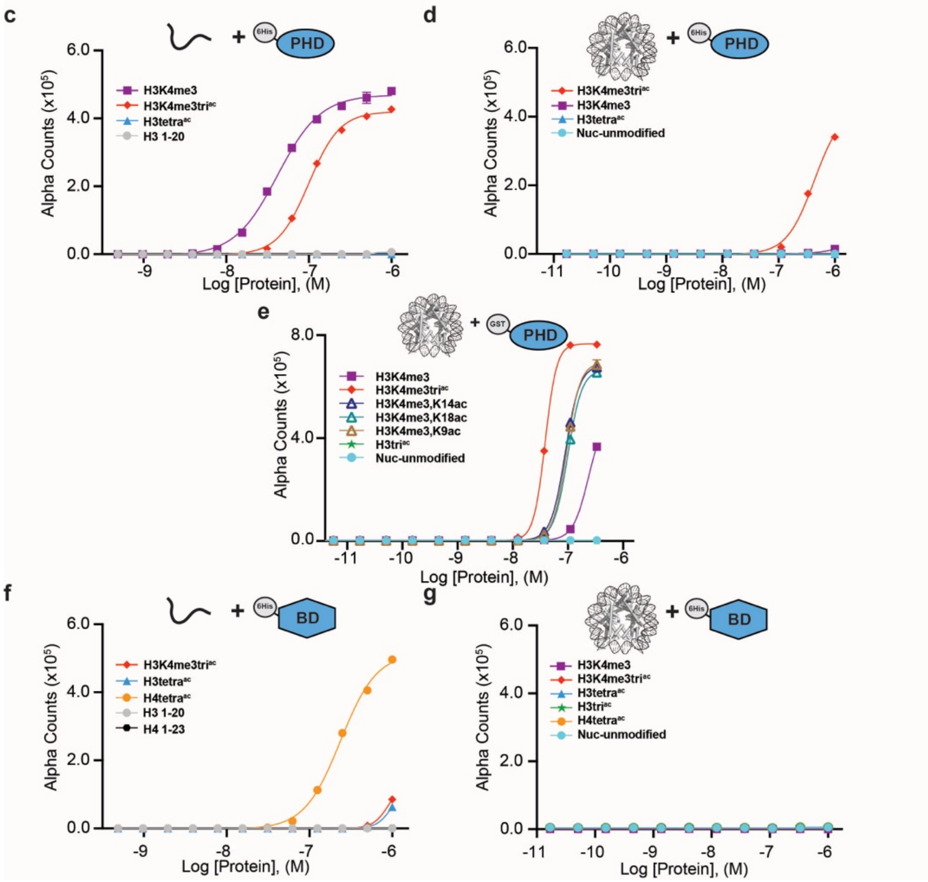


**\Extended Data Fig. 2. Individual BPTF reader domains have reduced affinity and restricted specificity in the Nuc context. a-b)** $EC_{50}^{rel}$ (with $CI^{95}$) values calculated from dCypher curves of GST-PHD, 6His-PHD, or 6His-BD Query binding to peptide (**a**) or Nuc (**b**) Targets from curves in (**c-g**), with same color code as per each legend. **c-e)** Alpha counts plotted as a function of 6His-PHD Query concentration for peptide (**c**) or Nuc (**d**) Targets, or for GST-PHD Query concentration for Nuc (**e**). **f,g)** Alpha counts plotted as a function of 6His-BD Query concentration for peptide (**f**) or Nuc (**g**) Targets. See previously[35] and **Suppl. Discussion** regarding comparison of $EC_{50}^{rel}$ across different affinity tags.

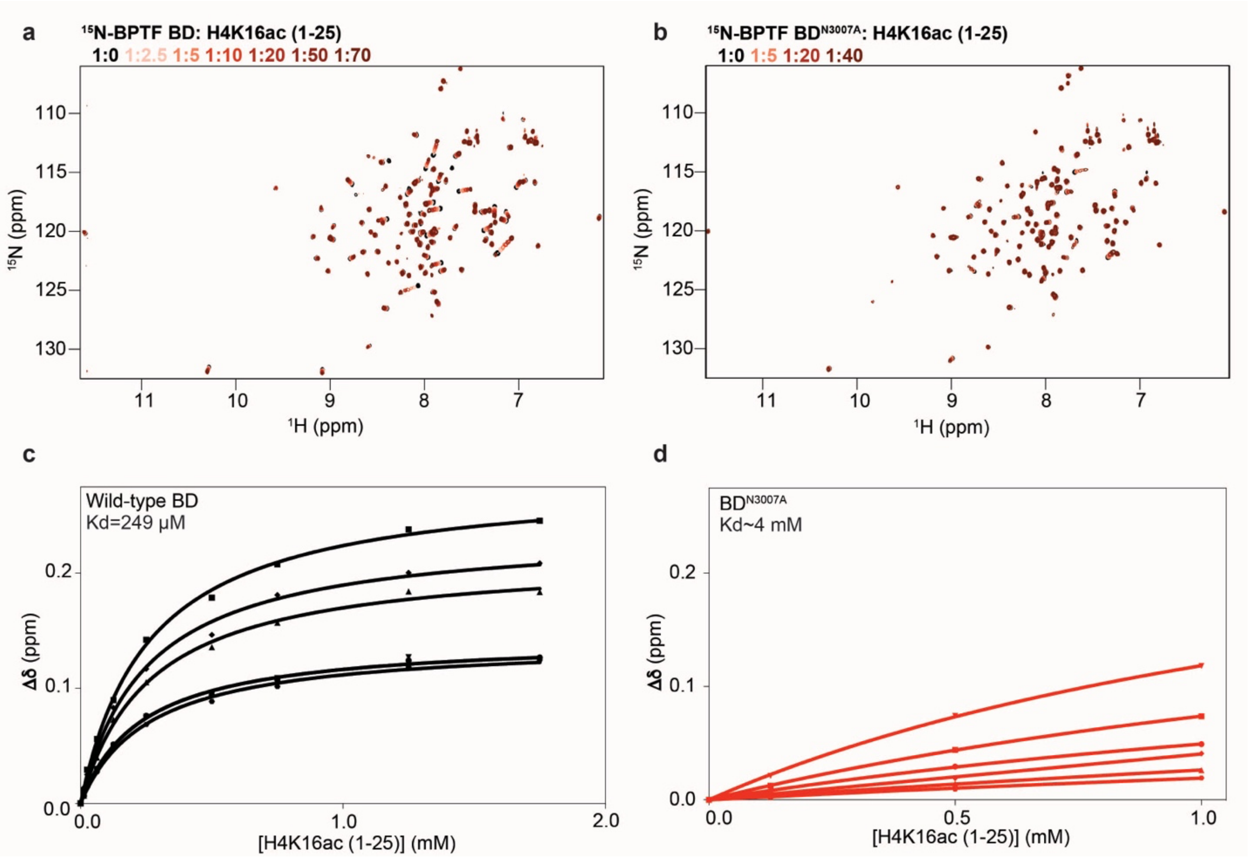


**Extended Data Fig. 3. NMR analysis of mutant BD. a-b)** Overlay of the $^{1}$H-$^{15}$N-HSQC spectra of $^{15}$N-BPTF BD wild-type (**a**) and $^{15}$N-BPTF BD-N3007A (BD$^{mut}$) (**b**) upon titration of H4K16ac (1-25) tail peptide **(Suppl. Tables 3: Resources D)**. Molar ratios are color coded as per legend. **c-d)** Binding curves for the five resonances with the highest chemical shift perturbations upon addition of peptide to $^{15}$N-BPTF BD wild-type (**c**) and $^{15}$N-BPTF BD-N3007A (BD$^{mut}$) (**d**). $K_d$ values were calculated using a single site binding isotherm accounting for ligand depletion (see **Methods**) and are denoted on each plot.

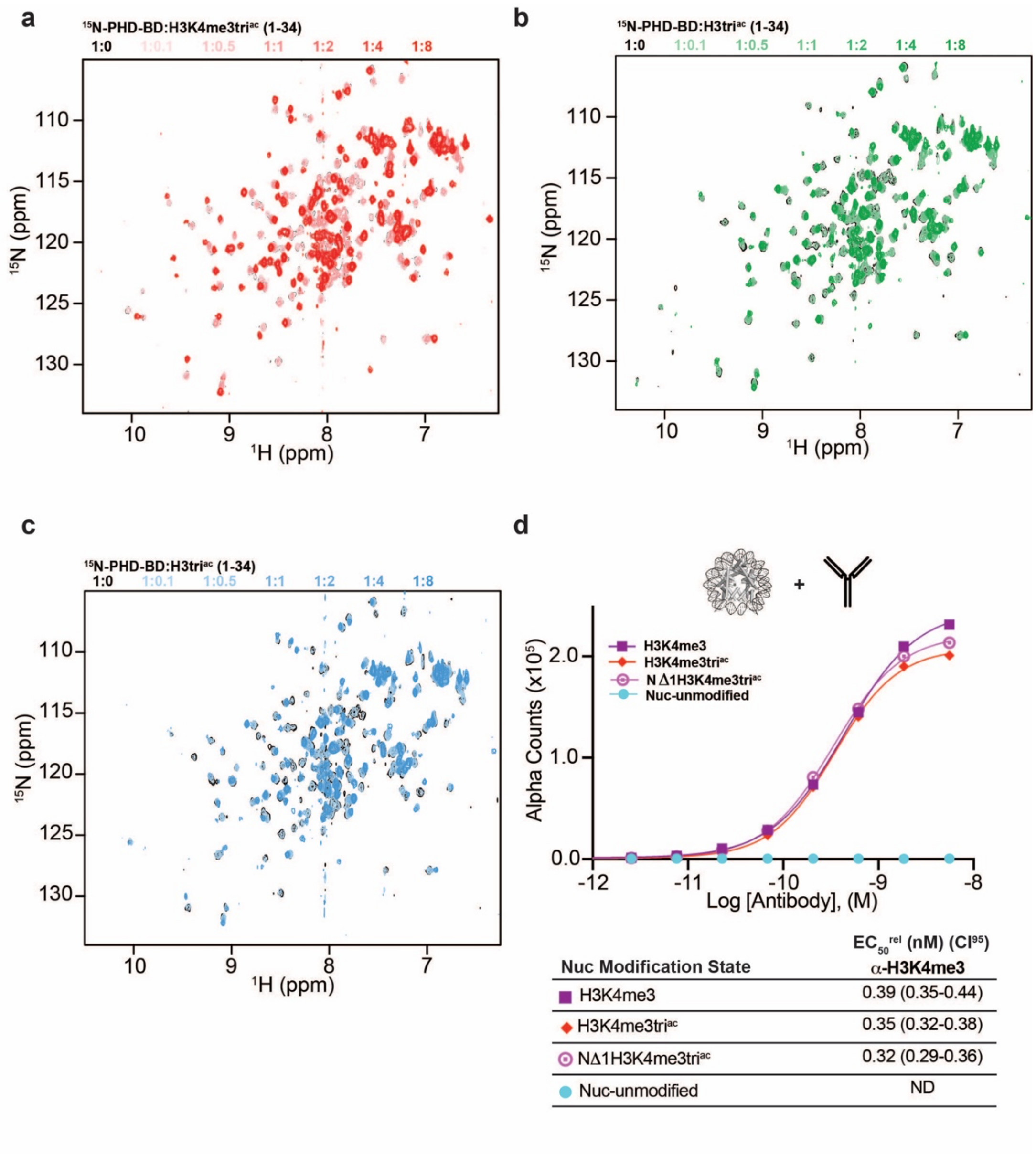


**Extended Data Fig. 4. Multivalent association with the H3 tail. a-c)** Full spectral overlays of $^{15}$N-PHD-BD on titration with histone H3 tail peptides (aa 1-34): H3K4me3tri$^{ac}$ (**a**); H3tetra$^{ac}$ (**b**); or H3tri$^{ac}$ (**c**). Molar ratios of each peptide are denoted by color in panel legends. **d)** Validation of truncated H3 Nucs (NΔ1H3K4me3tri$^{ac}$). Alpha counts plotted as a function of anti-H3K4me3 (*RevMab* Cat# 31-1039-00, lot # P-09-00676) concentration: binding of this reagent is dependent on H4K4me3 (see Nuc-unmodified) but independent of residue A1 (see NΔ1H3K4me3tri$^{ac}$) or acetylation at K9, K14 or K18. Table shows $EC_{50}^{rel}$ (with $CI^{95}$) values calculated from dCypher curves. Targets color coded as per legend.

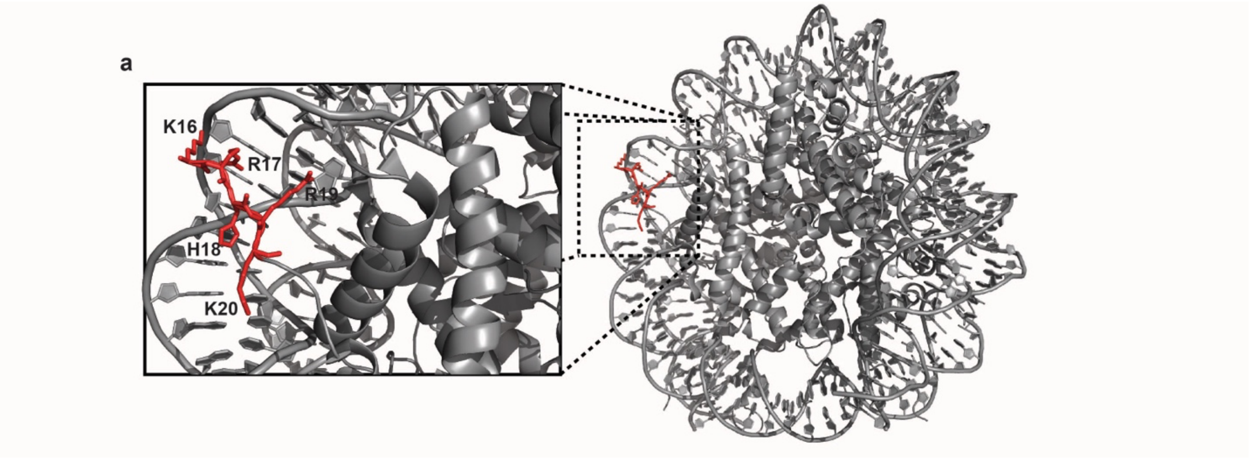


**Extended Data Fig. 5.** Nucleosome crystal structure (PDB ID: 3LZ0) showing interaction of the H4 tail basic patch (residues: K16-R17-H18-R19-K20) with DNA.

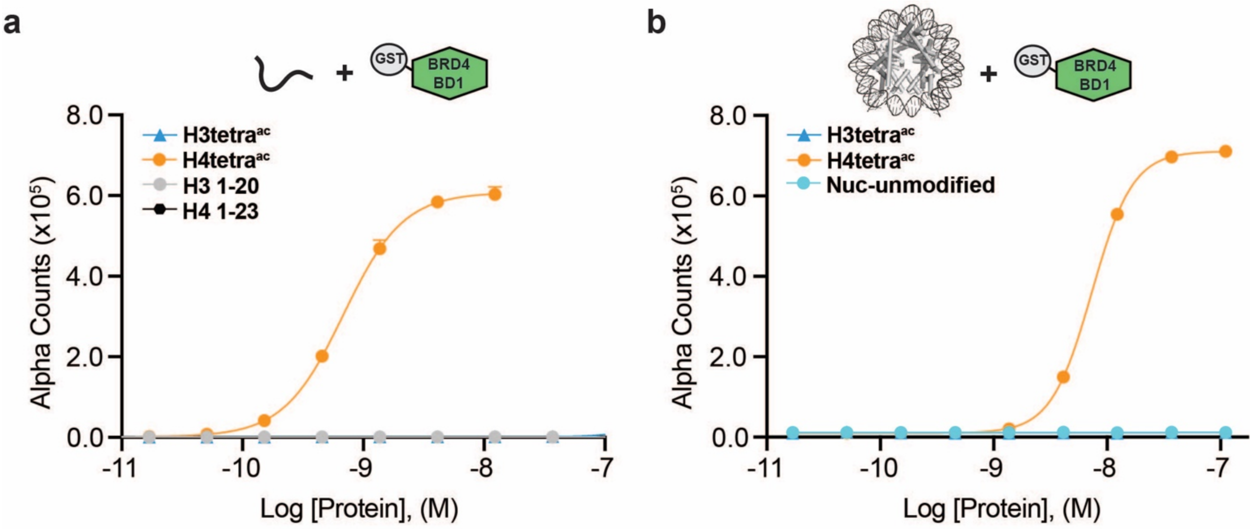


c

| Modification State | $EC_{50}^{rel}$ (nM) ($CI^{95}$) GST-BD1 Peptide | Nuc |
|---|---|---|
| ▲ H3tetra$^{ac}$ | >1000 | ND |
| ● H4tetra$^{ac}$ | 0.68 (0.64-0.73) | 7.3 (7.0-7.6) |
| ● NCP-unmodified | - | ND |
| ● H3 1-20 | ND | - |
| ● H4 1-23 | ND | - |

**Extended Data Fig. 6. GST-BRD4-BD1 binds acetylated histone H4 tail peptides and Nucs. a-b)** Alpha counts plotted as a function of GST-BRD4-BD1 Query concentration for peptide (**a**) or Nuc (**b**) Targets. **c)** $EC_{50}^{rel}$ (with $CI^{95}$) values calculated from dCypher curves in (**a**) and (**b**). Targets color coded as per legends. ND, Not Detected.

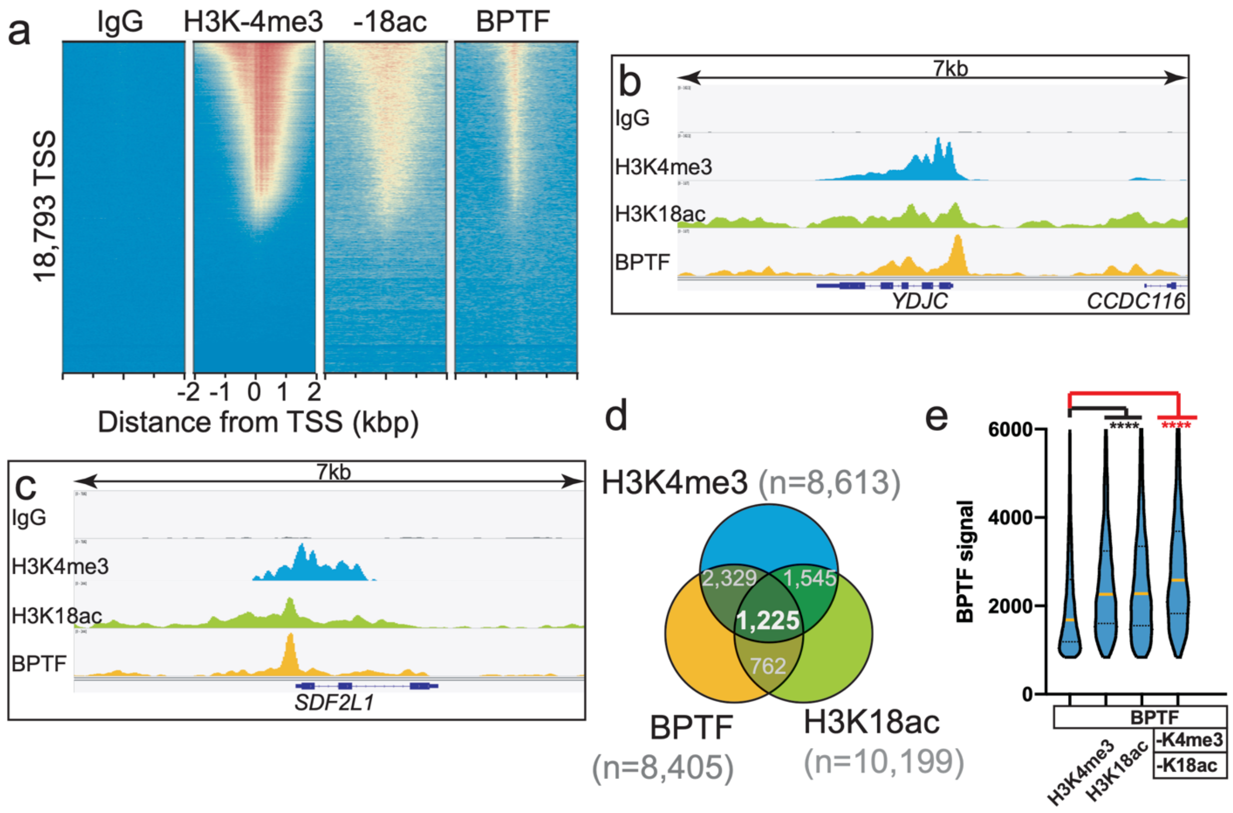


**Extended Data Fig. 7. The combinatorial PTM preference of BPTF PHD-BD *in vitro* is mirrored by its *in vivo* co-localization. a)** Heatmap of CUT&RUN signal (see **Methods**) aligned to the transcription start site (TSS, +/- 2kb) of 18,793 genes in K562 cells. High and low signal (red and blue respectively) are ranked by / linked to H3K4me3 (top to bottom). **b-c)** CUT&RUN RPKM normalized tracks at representative loci using Integrative Genomics Viewer (IGV, Broad Institute). Note the general co-localization of BTPF (endogenous) with H3K4me3 and H3K18ac. **d)** Venn overlap for H3K4me3, H3K18ac and BPTF peaks. Peaks were called using SEACR, designed for the Sparse Enrichment Analysis of CUT&RUN data[92], and intersected using bedtools[94]. To compare data sets, the top peaks (number noted for each) were considered. **e)** BPTF signal distribution for SEACR peaks represented as a Violin plot, and derived from Venn diagram groups. Column 1 depicts the signal distribution for all 8,405 BPTF peaks. Other columns depict BPTF peaks that overlap with peaks for H3K4me3, H3K18ac, or both PTMs (as noted). The Fisher exact test was used to determine if BPTF co-occupancy with specific PTMs or combinations were statistically significant (**** denotes P-value < $10^{-4}$). See **Suppl. Tables 3: Resources E** for sequencing statistics; all project files available at GEO accession GSE150617.

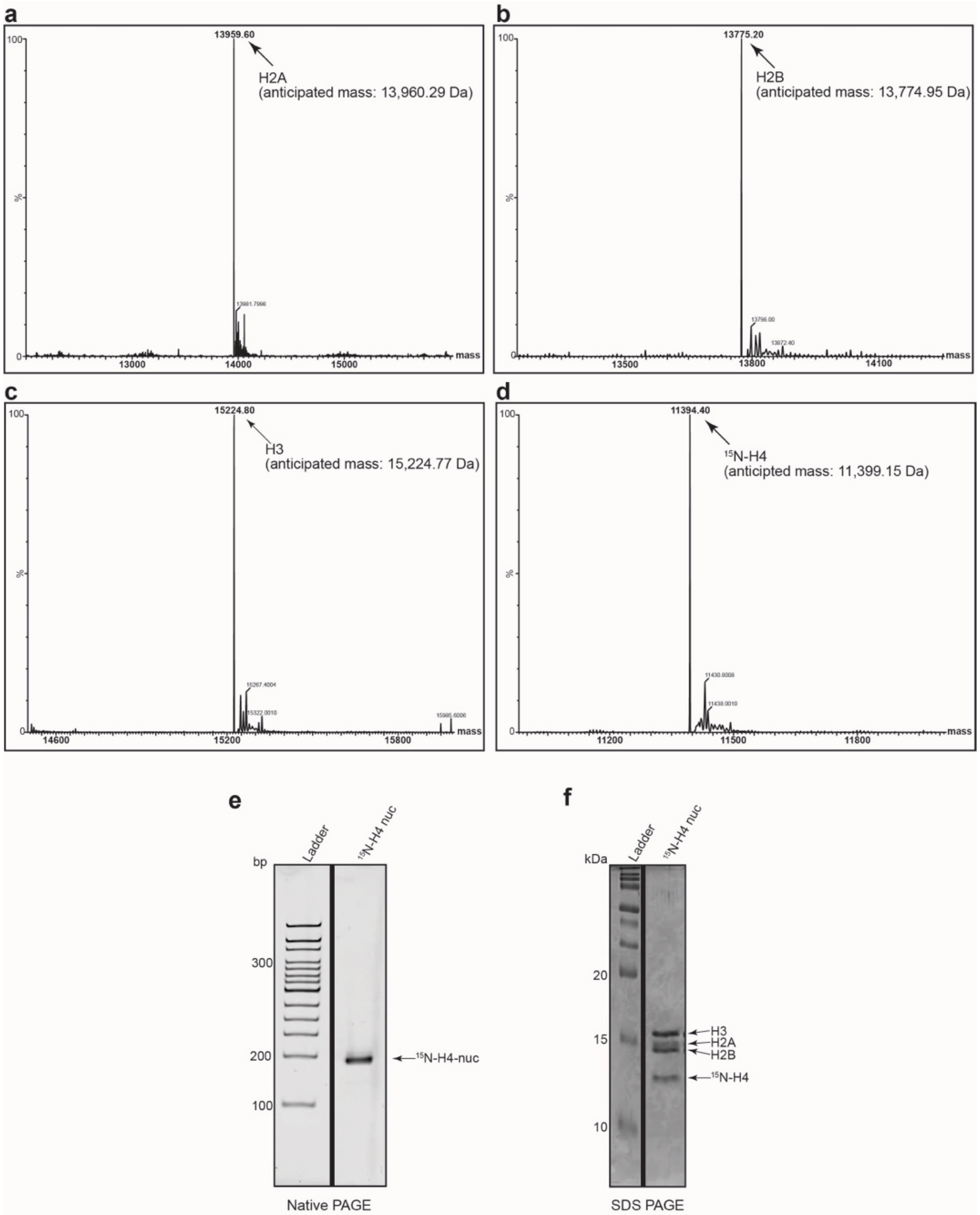


**Extended Data Fig. 8. Representative protein QC (NMR related). a-d)** Histones (including $^{15}$N-H4) for NMR were individually purified and validated by MALDI mass spectrometry prior to Nuc reconstitution (see **Methods**). **e)** The $^{15}$N-H4 containing Nuc was resolved by native PAGE and stained with ethidium bromide to demonstrate reconstitution / absence of free DNA. **f)** $^{15}$N-H4 containing Nucs were resolved by reducing SDS-PAGE and Coomassie stained to interrogate histone stoichiometry.

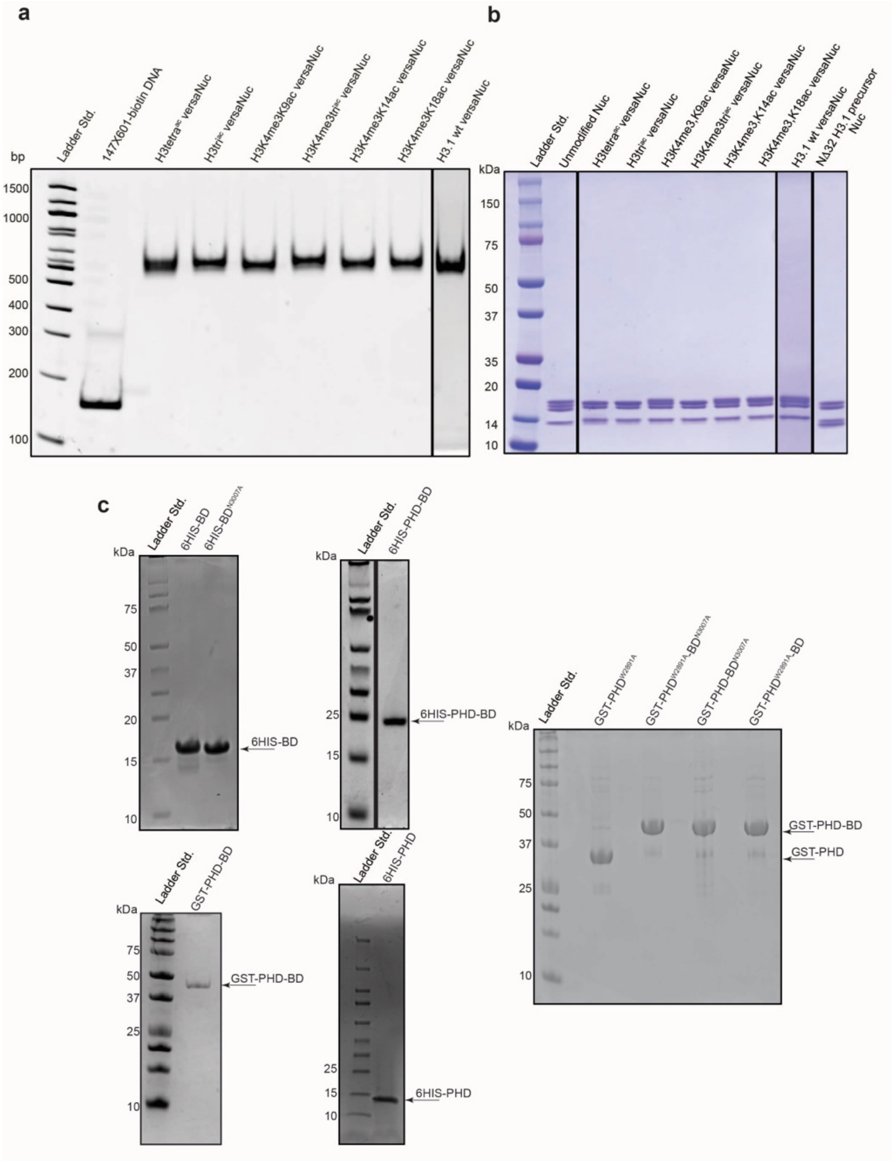

**Extended Data Fig. 9. Representative protein QC. a)** versaNucs (see **Suppl. Discussion**) (Lanes 1-7; PTM identity noted, 400 ng total DNA mass) were resolved by native PAGE (6%, 0.4X TBE) and integrity confirmed by staining with ethidium bromide (*vs.* Lane 8; 147X601 biotinylated free DNA, 200 ng total DNA mass). **b)** versaNucs (Lanes 1-7; 2 µg total protein mass) were resolved by reducing SDS-PAGE (4-20%) and histones visualized by Coomassie staining. Control lanes include unmodified Nuc (native full length H3; *EpiCypher* #16-6006) and versaNuc precursor (H3.1 NΔ32 before enzymatic ligation of H3 peptide; **Suppl. Tables 3: Resources D**). **c)** Indicated purified recombinants (see **Suppl. Tables 3: Resources A**) were resolved by SDS-PAGE and visualized by Coomassie staining.
